# Single-cell transcriptomes of developing and adult olfactory receptor neurons in *Drosophila*

**DOI:** 10.1101/2020.10.08.332130

**Authors:** Colleen N. McLaughlin, Maria Brbić, Qijing Xie, Tongchao Li, Felix Horns, Sai Saroja Kolluru, Justus M. Kebschull, David Vacek, Anthony Xie, Jiefu Li, Robert C. Jones, Jure Leskovec, Steven R. Quake, Liqun Luo, Hongjie Li

**Author notes:** co-first authors (Colleen N. McLaughlin, Maria Brbić).

## Abstract

Recognition of environmental cues is essential for the survival of all organisms. Precise transcriptional changes occur to enable the generation and function of the neural circuits underlying sensory perception. To gain insight into these changes, we generated single-cell transcriptomes of *Drosophila* olfactory receptor neurons (ORNs), thermosensory and hygrosensory neurons from the third antennal segment at an early developmental and adult stage. We discovered that ORNs maintain expression of the same olfactory receptors across development. Using these receptors and computational approaches, we matched transcriptomic clusters corresponding to anatomically and physiologically defined neuronal types across multiple developmental stages. Cell-type-specific transcriptomes, in part, reflected axon trajectory choices in early development and sensory modality in adults. Our analysis also uncovered type-specific and broadly expressed genes that could modulate adult sensory responses. Collectively, our data reveal important transcriptomic features of sensory neuron biology and provides a resource for future studies of their development and physiology.

## Introduction

Detection of sensory stimuli is critical for animals to find food, identify mates, recognize suitable habitats, and evade predators and harmful conditions. Specialized sensory neurons have evolved to discriminate and transmit information relating to chemical, thermal, and hygrosensory stimuli. In *Drosophila*, the ~50 types of primary sensory neurons that detect these cues are found in the third segment of the antenna and also the arista, a branched structure emanating from the antenna (Figure 1A). The majority of neurons in this sensory organ are olfactory receptor neurons (ORNs) that respond to a variety of volatile compounds (Hallem & Carlson, 2004; Hallem et al., 2006; Silbering et al., 2011). A subset of neurons in the antenna and arista respond to temperature and humidity-related stimuli (Yao et al., 2005; Barbagallo & Garrity, 2015; Enjin et al., 2016; Knecht et al., 2017). Each of the ~44 types of antennal ORNs expresses a distinct sensory receptor, or a unique combination of 2–3 receptors. Neurons that express the same receptor(s) also project their axons to the same glomerulus of the antennal lobe in the brain (Couto et al., 2005; Fishilevich & Vosshall, 2005; Benton et al., 2009; Silbering et al., 2011). Here, their axons form one-to-one connections with the dendrites of second-order projection neurons (PNs), thus creating discrete and anatomically stereotyped information processing channels (Figure 1A).

**Figure 1.**
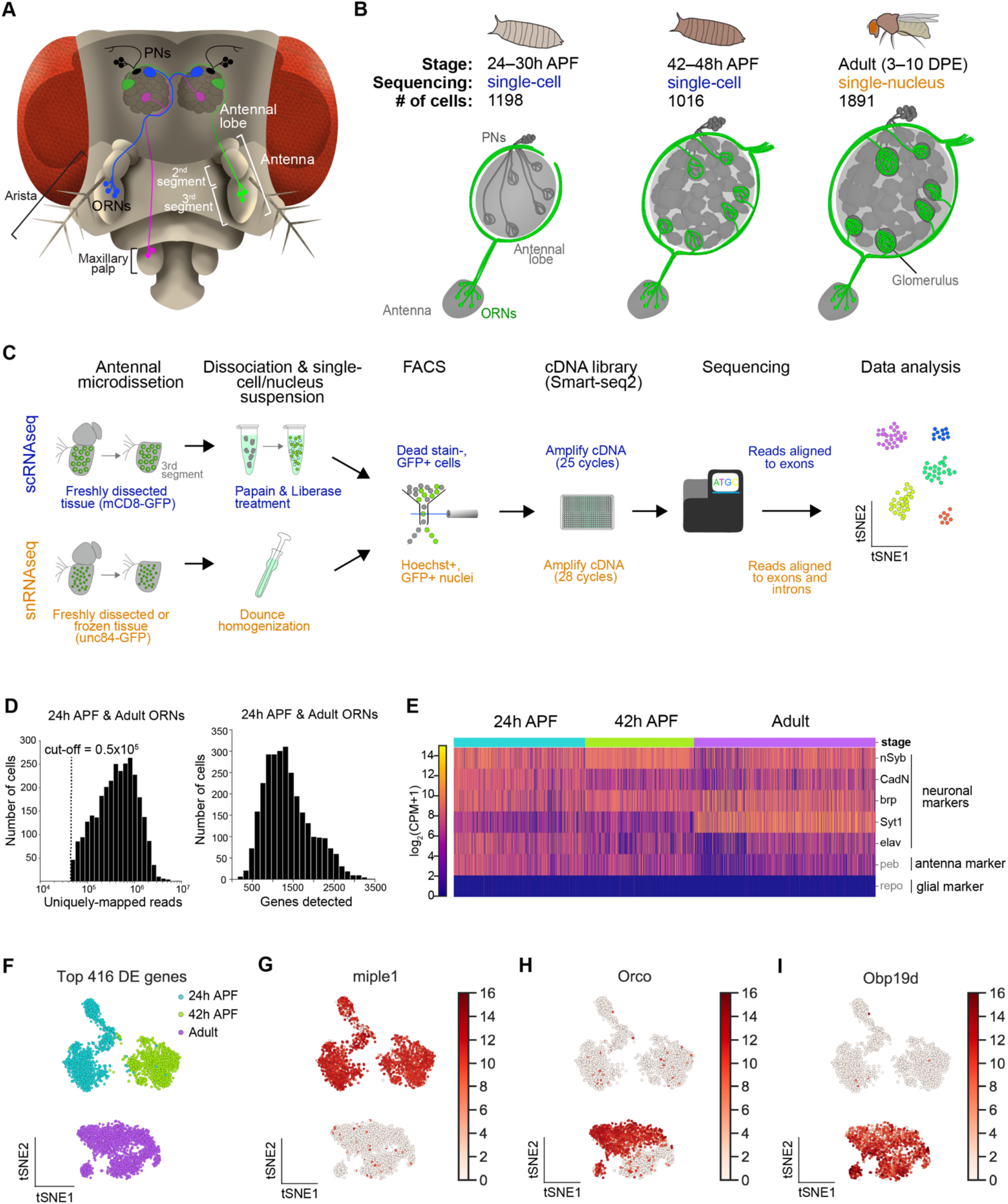
Single-cell transcriptomic profiling of *Drosophila* olfactory receptor neurons. (**A**) Schematic of *Drosophila* olfactory system. Three types of olfactory receptor neurons (ORNs) are portrayed in three different colors (green, magenta, and blue) and their cell bodies are housed in the antennae and maxillary palps. Axons form one-to-one synapses onto projection neuron (PN; black) dendrites in the antennal lobe of the brain. (**B**) Diagram containing information about the olfactory circuit and sequencing performed at three stages: 24–30h APF (h APF; hours after puparium formation), 42–48h APF, and adults (3–10 DPE; days post-eclosion). Each ORN type expresses a unique olfactory receptor or a unique combination of receptors and sends its axons to a stereotyped glomerulus. 42h APF neurons were previously sequenced in Li et al., 2020. (**C**) Diagram highlighting key differences in single-cell (blue) and single-nucleus (orange) RNA sequencing protocols. Cells are labeled with membrane-bound mCD8-GFP, whereas nuclei are marked with the nuclear envelop-bound unc84-GFP. (**D**) Distributions of uniquely mapped reads per cell (left) and number of detected genes per cell (right). Dashed line in uniquely mapped reads distribution marks the reads cut-off. (**E**) Heatmap depicting that ORNs analyzed express five pan-neuronal markers (*nSyb, CadN, brp, Syt1, elav*), an antennal cell marker (*peb*), but not a glial marker (*repo*). We filtered for high-quality neurons by ensuring that they expressed 2 out of 5 neuronal markers (in black) at log_2_(CPM+1) ≥ 2 (CPM: counts per million reads). (**F**) t-SNE plot depicting all sequenced ORNs. Dimensionality reduction was performed using the top 416 differentially expressed (DE) genes from neurons at all stages profiled. (**G–I**) Expression of genes specific to developing neurons (G) and mature neurons (H and I). Scale bar depicts log_2_(CPM+1).

Prior to carrying out their function in the adult, ORNs undergo multiple steps of development. This begins with their birth in the early pupal stage. Subsequently, ORNs extend their axons from the antenna into the brain where they take distinct trajectories around the antennal lobe, a process that positions them closer to their glomerular targets (Joo e al., 2013; Li et al., 2018) (Figure 1B, left). Axons next enter the antennal lobe to find and synapse with their partner PN’s dendrites (Jefferis et al., 2004; Mosca and Luo, 2014) (Figure 1B, middle). Concomitant with circuit formation and maturation, all ORN types express their unique olfactory receptor(s). Both the stereotyped development and organization of this system make it an excellent model to study molecular mechanisms of neuronal specification, axon guidance, and sensory physiology. However, little is known about the dynamic and cell-type specific transcriptional changes that occur to enable these distinct processes.

Single-cell RNA sequencing (scRNA-seq) is a powerful tool to investigate the mechanisms underlying nervous system development and function (Ofengeim et al., 2017; Mu et al., 2019; Li, 2020). Recent scRNA-seq studies in the *Drosophila* nervous system have enabled the discovery of new cell types, mechanisms important for development and aging, and genes controlling neural specification and wiring (Li et al., 2017; Croset et al., 2018; Davie et al., 2018; Avalos et al., 2019; Allen et al., 2020). Our recent study at a mid-developmental stage demonstrates that single-cell transcriptomic clusters represent anatomically and functionally distinct ORN types (Li et al., 2020). Yet, the mechanisms enabling sensory neuron types to transition through distinct developmental states into functional adult cells remain enigmatic. Furthermore, it has been difficult to obtain single-cell transcriptomic access to adult ORNs as these cells, like many other peripheral tissues in insects, are encapsulated within the hardened cuticle and cannot be isolated without substantial damage.

Here, we developed a single-nucleus RNA-seq (snRNA-seq) protocol to profile adult ORNs, thermosensory and hygrosensory neurons. We also used single-cell RNA-seq to profile these neurons at an early pupal stage. On a global level, we observed a shift in gene expression as neurons advance through development, from axon guidance-related transcripts to those important for sensory function. At the cell-type level, transcriptomic clusters corresponded to anatomically and functionally defined neuron types, allowing us to assign the cell-type identity of ~60% of developing and ~80% of adult clusters. Cell-type-specific transcriptomes, in part, reflected axon trajectory choice in early development but physiological functions in adults. Finally, we uncovered a set of genes in adult neurons that may function to regulate the detection or transmission of sensory information. Altogether, this dataset provides an important foundation for investigating molecular mechanisms controlling sensory neuron development and function.

## Results

### Single-cell transcriptomic profiling of *Drosophila* olfactory receptor neurons

Recently, we performed scRNA-seq on ORNs at a mid-developmental timepoint, 42–48h after puparium formation (APF; hereafter 42h APF), as their axons are in the final stage of selecting synaptic partner PNs (Li et al., 2020) (Figure 1B, middle). We sought to more completely understand how transcriptomic changes underlie the differentiation and function of these cells by profiling them at two additional stages: 24–30h APF and 3–10-day old adult (hereafter 24h APF and adult, respectively). For simplicity we will refer to the neurons within the third antennal segment as ORNs since they are the vast majority of neurons in this structure and their development and function are better studied, though a small number of hygrosensory and thermosensory neurons exist in this sensory organ (Barbagallo and Garrity, 2015) (see Figure 7). At 24h APF, ORN axons have just begun to circumnavigate the antennal lobe after selecting a specific trajectory, which is necessary for proper glomerular targeting (Figure 1B, left) (Joo et al., 2013). Thus, we reasoned that scRNA-seq of 24h APF neurons would provide us with a more comprehensive picture of their development and axon targeting. Adult single-cell transcriptomes are important for understanding how distinct ORN types detect and transmit sensory information, and adult cell types should be readily distinguishable by the specific sensory receptors they express. However, since adult ORNs are emmeshed in the hardened cuticle, they could not be readily isolated using standard scRNA-seq methods.

Therefore, we established a single-nucleus RNA-seq (snRNA-seq) protocol to profile adult ORNs (Figure 1C), as similar methods have been used to gain transcriptomic access to cell types that are resistant to standard dissociation protocols in other organisms (Habib et al., 2017; Denisenko et al., 2020; Ding et al., 2020; Kebschull et al., 2020). Our snRNA-seq protocol is comparable to our scRNA-seq method, in which GFP-labeled neurons in third antennal segment were dissected and dissociated into a single-cell suspension, sorted into 384-well plates via fluorescence activated cell sorting (FACS), and sequenced using the SMART-seq2 platform (Picelli et al., 2014) (Figure 1C, top). snRNA-seq differed from scRNA-seq at four steps (Figure 1C, bottom). First, instead of labeling neuronal membranes with *mCD8-GFP*, we labeled their nuclear envelopes with *unc84-GFP* (Henry et al., 2012). Second, nuclei were dissociated mechanically rather than enzymatically. Third, nuclei were labeled with a Hoechst, a fluorescent DNA stain, prior to being sorted in to 384-well plates via FACS. Finally, since nuclei contain less RNA content than whole cells, nuclear cDNA was amplified using 3 additional PCR cycles and sequencing reads were aligned to both exons and introns, instead of exons alone for scRNA-seq (Figure 1C; see Figure 2).

**Figure 2.**
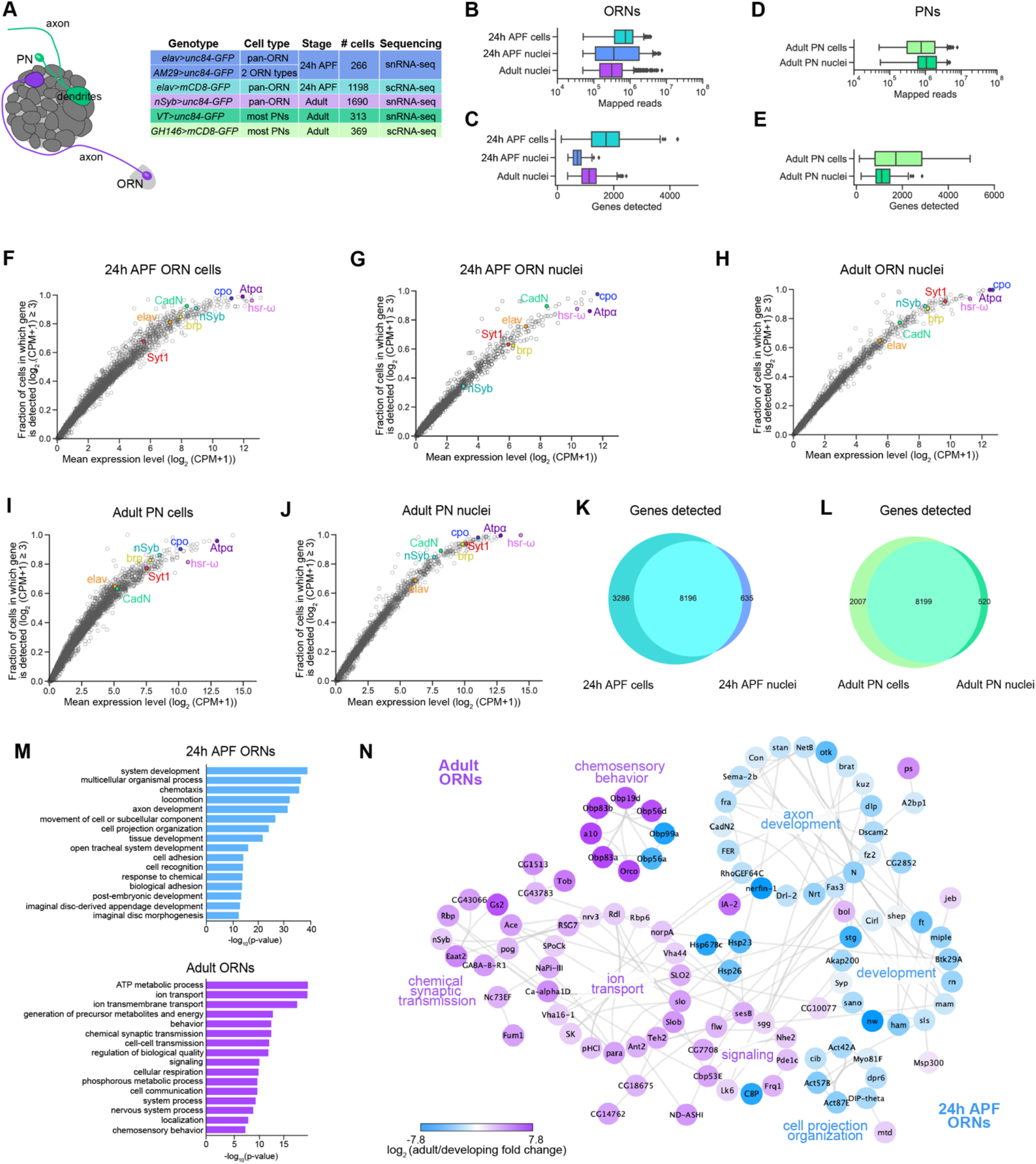
Comparison of single-cell and single-nucleus RNA-seq. (**A**) Schematic depicting ORN axons and PN dendrites in the antennal lobe (left) and summary table of the cell types, number of cells, genotypes, and stages that were profiled using snRNA-seq and scRNA-seq. Note that 24h APF cells were labeled by intersecting *elav-GAL4* with *ey-FLP* and *UAS-FRT-STOP-FRT-mCD8-GFP*. The reason there are fewer adult nuclei here than in Figure 1B is because some adult nuclei were labeled with lam-GFP, but those nuclei were not included in analyses here (see Figure 5—supplement 1C). (**B–E**) Box plots depicting uniquely-mapped reads (B, D) and detected genes (C, E) per cell/nucleus. Only exonic reads are shown for RNAs in cells, whereas exonic and intronic reads are combined for nuclear transcripts. (**F–J**) Mean expression level and detection rate of all genes in 24h APF ORNs (F), 24h APF ORN nuclei (G), adult ORN nuclei (H), adult PN cells (I), and adult PN nuclei (J). Detection is defined as log_2_(CPM+1) ≥ 3. Detection failure events can occur due to (1) the gene not being expressed in the cell; (2) gene dropouts resulting from technical artifacts even though mRNA transcripts are present; or (3) gene expression is below detection threshold. Colored circles represent the five neuronal markers used for quality control filtering (Figure 1E) and three house-keeping genes. (**K–L**) Venn diagram comparing all genes detected using scRNA-seq and snRNA-seq in 24h APF antennal neurons (J) and in adult PNs (K). Expression was defined as log_2_(CPM +1) ≥ 2. (**M**) Gene ontology (GO) analysis based on the top ~400 genes enriched in 24h ORN nuclei compared to adult ORN nuclei (top) and GO analysis of top ~400 genes enriched in adult ORN nuclei compared to 24h APF nuclei (bottom). Top GO 16 terms are shown. (**N**) STRING plot showing gene networks of 24h APF and adult ORN nuclei.

By combining scRNA-seq and snRNA-seq, we sequenced 24h APF and adult neurons to a depth of ~1 million reads per cell/nucleus. Following quality filtering for cells/nuclei with ≥ 50,000 reads that express at least 2 out of 5 neuronal markers (see Methods), we obtained 1,198 and 1,891 high-quality neurons at 24h APF and adult, respectively (Figure 1D, E). Together with the 1,016 42h APF ORNs (Li et al., 2020), we have a total of 4,105 neurons, representing three distinct developmental stages. Confirming transcriptomic quality, we observed high expression of neuronal markers, and the antennal cell marker *pebbled* (*peb*), but did not detect the glial marker *reversed polarity* (*repo*) in cells at all three stages (Figure 1E). Transcriptomes could be readily separated into their respective developmental stages following unbiased clustering using differentially expressed genes identified from all stages (Figure 1F). Further, we observed stage-specific expression of select genes, such as *miple1* in 24h and 42h APF ORNs, as well as adult-specific expression of the olfactory co-receptor *Orco* and the odorant binding protein *Obp19d* (Figure 1G–I). These data demonstrate the reliability of our single-cell and single-nucleus RNA-seq protocols and suggest that substantial transcriptomic changes occur as development proceeds.

### Comparison of single-nucleus and single-cell RNA-seq

Prior to examining ORN transcriptomes, we compared our snRNA-seq protocol to scRNA-seq. Since we were unable to directly evaluate adult ORN cells and nuclei, we sequenced 24h APF ORN nuclei to compare with cells at the same stage (Figure 2A). 24h APF ORN nuclei were labeled by driving nuclear-GFP with either the pan-neuronal *elav-GAL4* or *AM29-GAL4* which labels two ORN types (Figure 2A; Figure 2—supplement 1A) (Endo et al., 2007). We used pan-neuronal *nSyb-GAL4* to label adult ORNs. To evaluate if this technique is broadly applicable to fly neurons, we also sequenced adult PN nuclei the majority of which were labeled with *VT033006-GAL4* (Tirian & Dickson, 2017; see the companion manuscript by Xie et al. for details), to compare to stage-matched cells labeled with *GH146-GAL4* (Figure 2A).

We began by comparing the number of uniquely mapped reads and genes detected in cells to nuclei. We expected that snRNA-seq reads would be lower than scRNA-seq because nuclear transcripts are only a fraction of those found in the entire cell. When we aligned sequencing reads to exons only, both the number of mapped reads and genes detected in ORN nuclei were reduced compared to cells (Figure 2—supplement 1B, C). We tested if aligning nuclear sequencing reads to exons and introns could increase the gene detection rate, since intronic reads comprised 32.7% and 26.2% of total reads in 24h APF and adult ORN nuclei, respectively, compared to 10.5% when included in 24h APF cell reads (Figure 2—supplement 1D). In ORNs, we observed that inclusion of intronic reads increased average gene detection rate in 24h APF (from ~520 to ~700) and adult (from ~950 to ~1,100) nuclei, compared to 24h APF (~1,700) exonically aligned cell reads (Figure 2C, Figure 2—supplement 1C). Similarly, intronic reads were ~27% of total PN reads; when they were added into exonic reads, the mean number of genes detected increases from ~1,000 to ~1,200 in nuclei (Figure 2D, E; Figure 2—supplement 1E). Thus, including intronic reads in snRNA-seq data provides additional information that is not captured by exonic reads alone.

We next assessed gene expression variability between cells and nuclei. Changes in gene expression between cells can arise from biological factors, such as differences in transcription rate or cell state, or are the result of technical artifacts leading to missed gene detection or “gene dropouts” (Li & Xie, 2011; Munsky et al., 2012; Little et al., 2013). We evaluated expression of the five neuronal markers used for quality filtering of cells and nuclei (*elav, CadN, Syt1, brp, nSyb*) and house-keeping genes (*cpo, Atpa, hsr-ω*) to determine the contribution of gene dropouts to expression variability. The majority of genes we evaluated showed similar expression in both ORN and PN cells and nuclei (Figure 2F–J). However, we did notice that some genes, namely *nSyb* and *brp*, had a higher dropout rate in 24h nuclei compared to cells at the same stage (Figure 2F, G), a finding that is in line with a previous report in mammalian neurons (Bakken et al., 2018). These results indicate that gene dropouts likely do not contribute to considerable expression differences between cell and nuclear transcriptomes.

Next, we evaluated the extent to which snRNA-seq and scRNA-seq can detect similar genes. We first assayed the 20 highest expressed genes by percentage of total reads and found that stage-matched cells and nuclei share a subset of the same highly expressed genes (Figure 2—supplement 1F vs. G, I vs. J). As expected, we found that ORN and PN cells show higher detection of mitochondrial transcripts than nuclei, with the exception of adult ORN nuclei which express some mitochondrial transcripts likely resulting from dissociation artifacts (Figure 2—supplement 1H). Second, we compared the total genes detected in cells and nuclei. We observe that 71% of genes detected in ORN cells were found in nuclei, and 93% of nuclear genes were detected in cells (Figure 2K). Likewise, 80% of genes detected in PN cells were detected in their nuclear counterparts, and 94% of genes detected in PN nuclei were in cells (Figure 2L), indicating that snRNA-seq can detect similar genes to scRNA-seq. Finally, we queried the types of genes that were differentially expressed between cells and nuclei by performing gene ontology (GO) analysis between stage-matched cell and nuclear transcriptomes (The Gene Ontology Consortium, 2000). Based on studies in other species (Bakken et al., 2018; Deeke & Gagnon-Bartsch, 2020), we predicted that cells would be enriched for genes with house-keeping functions, probably resulting from preferential cytoplasmic mRNA stability. Indeed, cellular transcriptomes were enriched for genes in GO categories such as translation, protein folding, and metabolic processes, whereas nuclear transcriptomes contained terms related to stage-specific processes, e.g., neuron differentiation, when compared to cells (Figure 2—supplement 1K, L). Most of the stage-specific genes were also expressed in ORN cells, albeit at slightly reduced levels relative to nuclei (Figure 2—supplement 1N, last column), indicating that our snRNA-seq and scRNA-seq protocols detect mostly overlapping genes. RNAs expressed from the mitochondrial genome were enriched in ORN cells, whereas long non-coding RNAs were enriched in nuclei (Figure 2—supplement 1M, N).

Finally, we asked if snRNA-seq could reveal transcriptomic changes associated with neuronal maturation. Thus, we computed the differentially expressed genes between nuclei at 24h APF and the adult stage and identified GO terms for each timepoint. 24h APF transcriptomes were enriched for terms that describe axon and tissue development, cell adhesion, and cell recognition (Figure 2M, N). Adult specific-genes, on the other hand, were involved in processes like signaling, cell communication, and chemosensory behavior (Figure 2M, N). These results indicate that as ORNs mature, there is a switch in expression from genes important for axon guidance and circuit formation to genes that mediate neuronal signaling and odorant detection. Altogether, our snRNA-seq protocol can robustly and reliably profile developing and adult neurons.

### Individual ORNs maintain the expression of the same olfactory receptor(s) throughout development

ORNs use three distinct families of olfactory receptors to detect odorants: Ors (odorant receptors), Irs (ionotropic receptors), and Grs (gustatory receptors) (Clyne et al., 1999; Vosshall et al., 1999; Scott et al., 2001; Benton et al., 2009). Olfactory receptor genes are gradually turned on in the pupal stage and by the time they reach adulthood each neuron type expresses one or few specific receptor(s) (Figure 3A). Olfactory receptors can, therefore, serve as excellent ORN type-specific markers. In the mouse olfactory system, an immature ORN can express several olfactory receptors before selecting the one whose expression will be maintained in the mature cell (Hanchate et al., 2015; Tan et al., 2015). Olfactory receptor expression has not been thoroughly evaluated in *Drosophila* ORNs during development. Thus, we examined this issue more closely.

**Figure 3.**
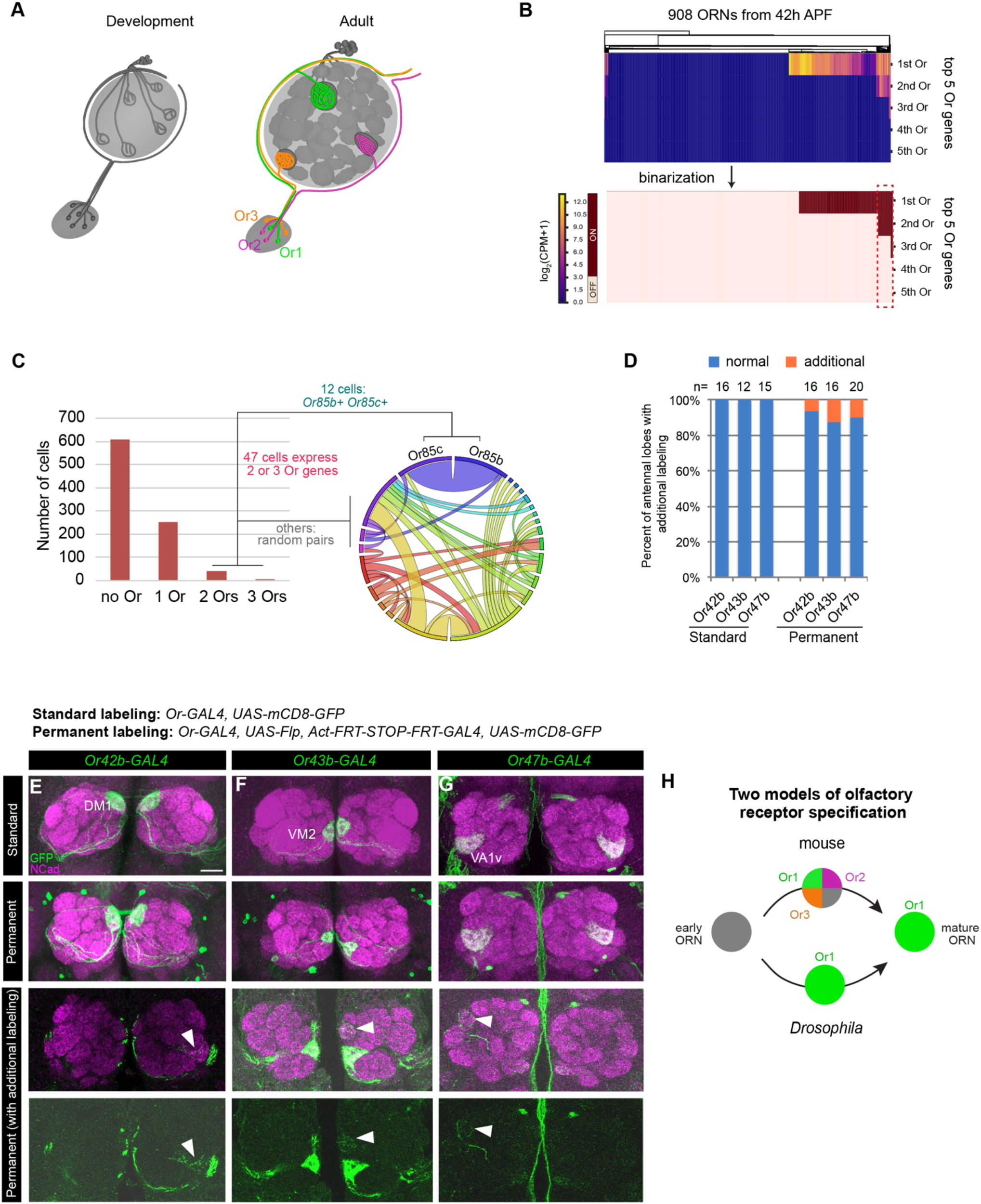
Individual ORNs maintain the expression of the same olfactory receptor(s) throughout development. (**A**) Schematic depicting ORNs at an early developmental stage (left) and in the mature circuit (right) where each ORN type expresses a distinct olfactory receptor represented by different colors. (**B**) Heatmap (top) and binarization of heatmap (bottom) of the top 5 highest expressed olfactory receptors in 908 *nSybGAL4>mCD8-GFP* positive ORNs from 42h APF. Olfactory receptors were considered “on” if they were expressed at log_2_(CPM+1) ≥ 3. (**C**) Quantification of the number of cells expressing zero, one, two, or three olfactory receptors in 42h APF ORNs (left). Chord plot (right) of co-expressed receptors in the 47 cells that expressed 2 or 3 olfactory receptors. Most co-expressed receptors show random patterns. (**D**) Quantification of the percentage of antennal lobes that displayed labeling of additional glomeruli in standard and permanent labeling conditions. (**E–G**) Confocal images of adult antennal lobes labeled with anti-GFP (green) and anti-NCad (magenta) in standard labelling (top row) or permanent labeling (bottom three rows) conditions of *Or42b-GAL4-positive* (E panels), *Or43b-GAL4-positive* (F panels), *Or47b-GAL4-positive* (G panels) ORNs. Antennal lobes that showed labeling of additional glomeruli are depicted in the last two rows. Arrowheads denote additional labeling. Scale bar, 20 μm. (**H**) Schematic depicting olfactory receptor specification in mice and *Drosophila*. Mouse ORNs express multiple olfactory receptors before one predominates in mature ORNs (top; Hanchate et al., 2015; Tan et al., 2015), whereas our data indicate that *Drosophila* ORNs select a single olfactory receptor to express during development that is maintained in adulthood (bottom).

Our scRNA-seq data indicated that some ORNs began to express Ors at 24h APF (Figure 4C), and by 42h APF roughly one-third of ORNs express these receptors (Figure 3B, C). Of the 298 Or-expressing ORNs at 42h APF, 84% (251 cells) express only one receptor and 16% (47 cells) express two or three (Figure 3B, C). Some ORN types are known to express more than one Or (e.g., Or19a/b and Or65a/b/c). Prior to this study *Or85c* was thought to be expressed in the larval olfactory system (Kreher et al., 2005), but we detect co-expression of *Or85c* with *Or85b* in 4% (12 cells) of *Or*-expressing ORNs (Figure 3C). All other co-expressed Ors appeared to be randomly paired (Figure 3C). Low-frequency co-expression could be accidental or be the result of doublet sequencing, an infrequent event whereby the transcriptomes of two cells are inadvertently combined during sample processing or sequencing (Ilicic et al., 2016). We conclude that the majority of developing ORNs express only one Or gene.

**Figure 4.**
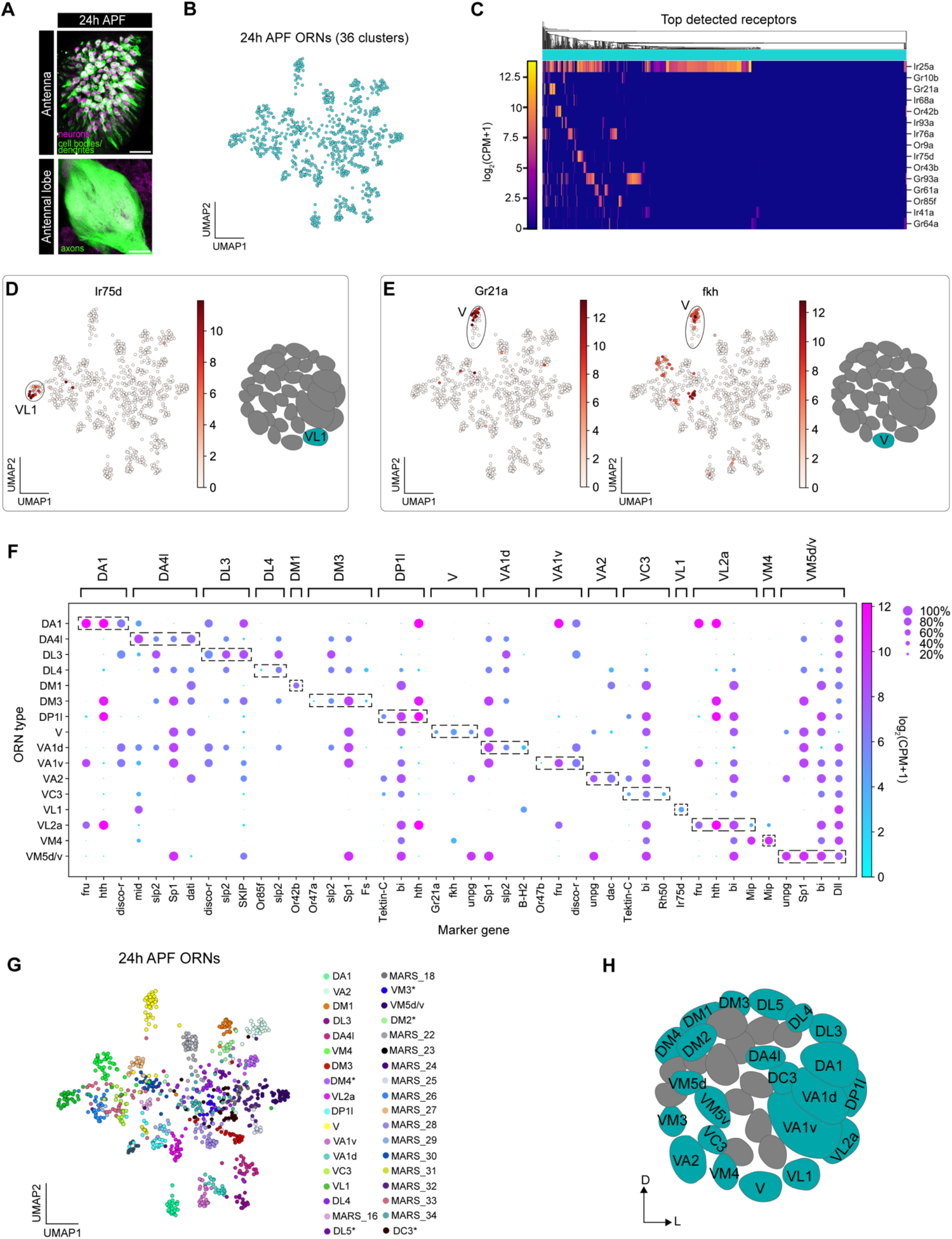
Mapping 24h APF transcriptomic clusters to their glomerular types. (**A**) Confocal images of 24h APF *elav-GAL4*, *ey-FLP*, *UAS-FRT-STOP-FRT-mCD8-GFP* neurons in the antenna (top) labeled with anti-GFP (green) and anti-Elav (magenta) and antennal lobe (bottom) labeled with anti-GFP (green). Scale bar, 20 μm. (**B**) UMAP plot depicting 36 clusters of 24h APF ORNs. Only *acj6*-positive neurons were used for this analysis. Dimensionality reduction was performed using differentially expressed genes from 42h APF combined with olfactory receptors for a total of 335 genes. (**C**) Heatmap depicting the top detected olfactory receptors in 24h APF neurons; detection threshold was defined as log_2_(CPM+1) ≥ 3 in > 5 neurons. (**D**) UMAP plot depicting *Ir75d* expression in a single cluster (left). *Ir75d-positive* ORNs target their axons to the VL1 glomerulus as shown in the schematic (right). (**E**) UMAP plots (left) depicting *Gr21a* expression in a single cluster and *fkh* expression in the same cluster. *Gr21a*-positive neurons target their axons to the V glomerulus and *fkh* is a marker expressed in V neurons at 24h and 42h APF. Schematic of glomerular projection of V neurons (right). (**F**) Dot plot summarizing the markers used to map 17 transcriptomic clusters at 24h APF to their corresponding 42h APF clusters. Each dot represents: mean expression within each cluster (color) and fraction of cells within the cluster expressing the gene (dot size). Dashed boxes around dots highlight markers used to map 24h APF clusters to 42h APF ones. (**G**) UMAP plot depicting the 22 transcriptomic clusters that were decoded at 24h APF. Asterisk next to ORN type indicates that it was annotated using data presented in Figure 6. Note that VM5d/v could not be separated (see Figure 4—supplement 2). The identity of clusters marked “MARS_#” is unknown. (**H**) Schematic depicting the glomerular targeting patterns of the 22 transcriptomic clusters mapped in (F–G; and Figure 6). Heatmap scale bars are in log_2_(CPM+1).

Do ORNs maintain expression of the same Or from the pupal to adult stage? To address this question, we employed a permanent genetic labeling strategy, which allowed us to label the membrane and trace the glomerular projection of any ORN that has expressed a specific *Or-GAL4* at any time. Specifically, we used an *Or-GAL4* to drive *UAS-FLP* to excise the STOP cassette between the ubiquitous *actin5C* promoter and *GAL4* transcriptional activator, such that any cell that transiently expresses the *Or-GAL4* will be labeled. *Or-GAL4* driving expression of *UAS-mCD8-GFP* served as the “standard labeling” control. We tested *GAL4* lines of three highly expressed Ors at 42h APF: *Or42b, Or43b*, and *Or47b* using both labeling methods (Figure 3—supplement 1A). In the majority of cases, permanently labeled ORN axons projected to the same glomerulus as the standard labeled controls (Figure 3D–G, Figure 3—supplement 1B–D’), indicating that these Ors were not expressed in multiple ORN types during development. Therefore, *Drosophila* and mouse ORNs appear to use different strategies for selecting specific olfactory receptor expression (Figure 3H).

The finding that olfactory receptor expression is stably maintained across development in *Drosophila* enabled us to use cluster-specific receptor expression to match transcriptomic clusters of developing ORNs to their respective ORN types, which also corresponds to the glomerulus their axons innervate (Couto et al., 2005; Fishilevich & Vosshall, 2005; Silbering et al., 2011).

### Mapping 24h APF transcriptomic clusters to their glomerular types

We next investigated the individual features of ORN transcriptomes at the two newly profiled stages. We started with 24h APF. When we visualized 24h APF transcriptomes in the t-SNE space (van der Maaten & Hinton, 2008) following dimensionality reduction with 500 over-dispersed genes from this stage, 24h APF transcriptomes separated into three distinct groups instead of forming clusters corresponding to each neuron type (Figure 4—supplement 1A). Upon querying expression of transcripts known to be expressed in ORNs such as *acj6*, a transcription factor that is ubiquitously expressed at 42h APF (Li et al., 2020), we found that *acj6* was enriched in the largest cluster but was absent from the other two clusters (Figure 4—supplement 1B). We asked if Acj6 protein reflected its transcript expression in the 24h APF antenna by co-immunolabeling with Acj6 and the pan-neuronal marker Elav. Given Acj6’s role in ORN axon targeting and olfactory receptor expression (Clyne et al., 1999a; Komiyama et al., 2004; Li et al., 2020) it could be turned on at a later stage of ORN maturation than Elav. Indeed, some neurons in the third antennal segment were Acj6-negative, and Acj6 was absent from the second antennal segment, which contains auditory neurons (Figure 4—supplement 1C). We assayed if either of the *acj6*-negative clusters could represent auditory neurons. Accordingly, the small *acj6*-negative cluster contained hearing-related genes (e.g., *iav*) (Senthilan et al., 2012) and GO terms, indicating that this cluster contains auditory neurons (Figure 4—supplement 1B, D–F). We performed identical analyses on the large cluster of acj6-negative cells and found that it contained genes/GO terms relating to neurogenesis, cell component biogenesis, and sensory organ development, whereas neurite development and cell signaling were found in *acj6*-positive cells (Figure 4—supplement 1F, G). Although all ORNs are born during the pupal stage, the precise time when their generation ceases is unknown (Endo et al., 2007; Barish & Volkan, 2015; Li et al., 2018). Thus, it is possible that we captured two distinct developmental states of ORNs with the large *acj6*-negative cluster being less mature than the *acj6*-positive cells.

Next, we sought to map 24h APF transcriptomes to their respective glomerular types (Figure 4A). We focused on the putatively more mature *acj6*-positive ORNs and began by identifying a gene set for dimensionality reduction that would enable us to obtain high-resolution separation of these cells. In the companion manuscript that analyzed single-cell transcriptomes of PNs (Xie et al.), we showed that using differentially expressed genes from the stage with maximal transcriptional diversity between different cell types for dimensionality reduction achieved the best clustering results. For the three stages of ORNs that we profiled, we found that transcriptomes at 42h APF exhibited highest diversity (Figure 4—supplement 2A). We therefore used differentially expressed genes from 42h APF ORNs in combination with known antennal olfactory receptors (Grabe et al., 2016) as the gene set for dimensionality reduction, and obtained clear separation of the *acj6*-positive 24h APF transcriptomes (hereafter 24h APF ORNs) when embedded in the UMAP space (McInnes et al., 2018) (Figure 4B). Neither sequencing batch nor sequencing depth drove clustering (Figure 4—supplement 2B, C).

We used the recently developed method, meta-learned representations for single cell (MARS) to cluster 24h APF ORNs (Brbic et al., 2020). MARS groups cells according to their cell types using a deep neural network to learn a set of cell-type specific landmarks and embedding function to project cells into a latent low-dimensional space. MARS segregates 24h APF transcriptomes into 36 distinct clusters (Figure 4—supplement 2D), which corresponded well with clusters identified by the density-based HDBSCAN approach (Campello et al., 2013). However, MARS separated closely related clusters that represent multiple ORN types better than HDBSCAN (Figure 4—supplement 2D, E). In fact, MARS was able to separate two clusters, VM5d and VM5v (Figure 4—supplement 2F), in our 42h APF data that were previously indistinguishable (Li et al., 2020). Thus, we used MARS to cluster and annotate 24h APF ORNs.

We first asked if we could detect expression of any olfactory receptors at 24h APF, as receptor expression in a single cluster would enable us to immediately identify the glomerular type the cluster represents. We observed expression of a small subset of receptors at 24h APF (Figure 4C). Cluster-specific expression of six of these receptors allowed us to annotate clusters corresponding to V (*Gr21a*), DM1 (*Or42b*), VL1 (*Ir75d*), DL4 (*Or85f*), VA1v (*Or47b*), and DM3 (*Or47a*) neuron types (Figure 4D–G, Figure 4—supplement 3A, B, D).

We next used marker genes to match additional 24h APF clusters with their corresponding 42h APF clusters. To annotate 42h APF clusters, we identified cluster-specific markers and used *GAL4* lines for each gene to validate that neurons expressing each marker targeted in the correctly annotated glomerulus (Li et al., 2020). Since axons at 24h APF have not reached their final glomerular positions, we could not perform similar analyses at this stage. Instead, we asked if cluster-specific markers from 42h APF displayed comparable expression at 24h APF and could be used to annotate clusters at this earlier stage. Demonstrating the feasibility of this strategy, *fkh* was expressed in neurons projecting to the V glomerulus at both 24h and 42h APF (Figure 4E, F, Figure 4—supplement 2F) (Li et al., 2020). We identified a set of 19 additional marker genes whose individual or combinatorial expression was sufficient to map ORNs at 24h APF to their respective 42h APF clusters (Figure 4F, Figure 4–supplement 2F). Using this approach, we decoded the glomerular identity of an additional 11 ORN types (Figure 4G, H, Figure 4—supplement 3). Two of these ORN types, VM5d and VM5v, mapped to a single cluster at 24h APF (Figure 4G), indicating that these cells are highly similar at this stage. Collectively, we mapped 16 transcriptomic clusters to 17 neuron types at 24h APF using olfactory receptor and marker gene expression. Additional clusters were mapped to neuronal types by matching transcriptomic clusters across development (see Figure 6).

### Mapping adult transcriptomic clusters to their glomerular types

The adult antenna is covered in hair-like projections called olfactory sensilla and has a large feather-like protrusion known as the arista (Figure 5A). Individual antennal ORN types are housed in one of four sensillar classes (see Figure 7A). Each individual sensillum comprises 1–4 ORNs and a few accessory cells that are all derived from a common progenitor (Figure 5A’) (Endo et al., 2012). The arista contains sensory neurons that express Irs and Grs and project their axons into the antennal lobe (Ni et al., 2013; Enjin et al., 2016). Unlike some pupal sensory neurons, all adult ORNs should express their type-specific receptor(s), enabling us to definitively map adult clusters to their glomerular type (Figure 5B). We observed that 75% of antennal sensory receptors are highly expressed in adult transcriptomes and 84% of adult neurons express at least one sensory receptor (Figure 5C). Dropouts and low-level receptor expression likely account for the 16% of neurons where we could not detect a receptor.

**Figure 5.**
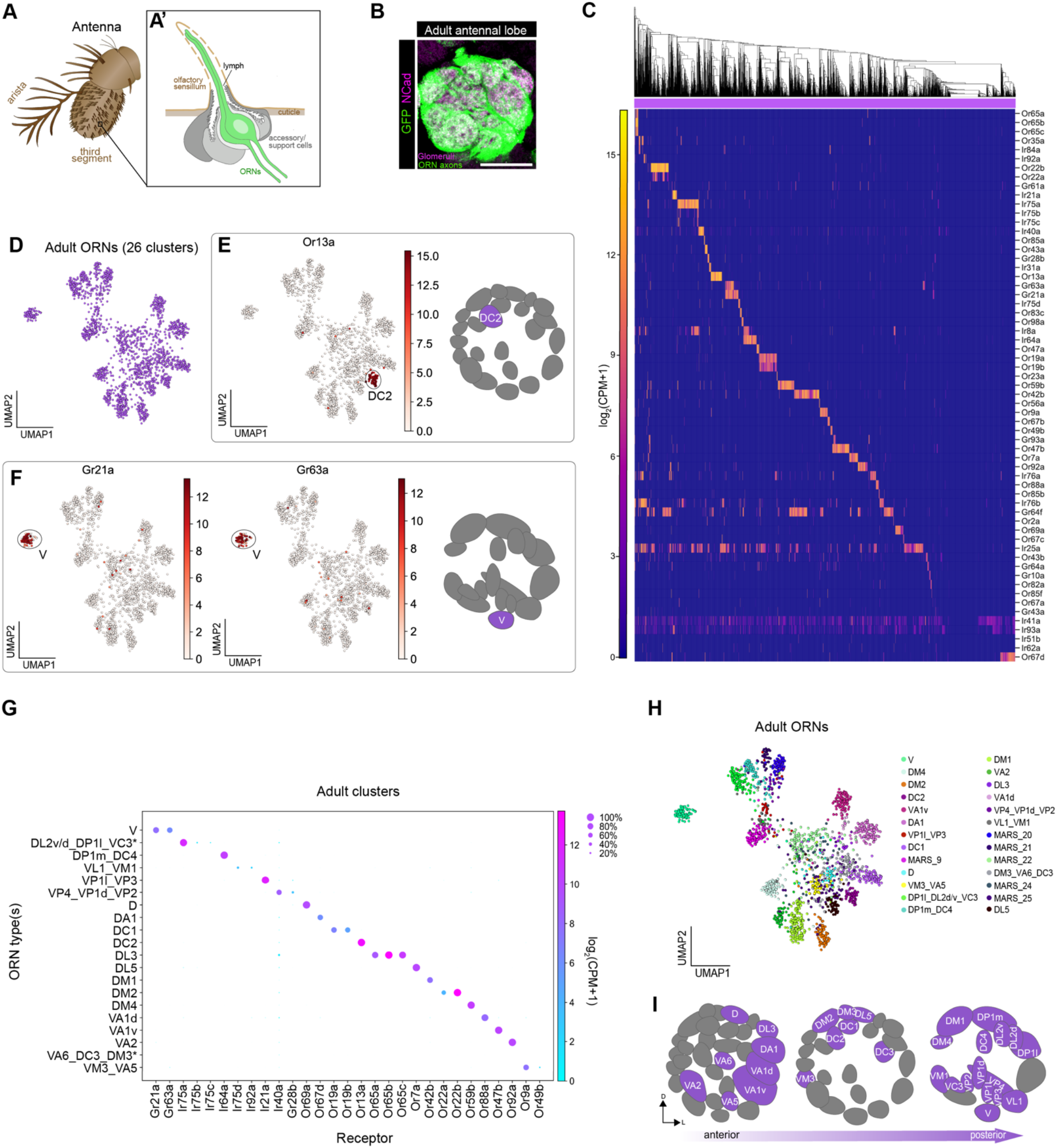
Mapping adult transcriptomic clusters to their glomerular types. (**A–A’**) Diagram of the adult antenna (A) and the structure of an individual sensillum (A’) where ORN dendrites and support cells are located. (**B**) Confocal image of *nSyb-GAL4; ey-FLP; UAS-FRT-STOP-FRT-mCD8-GFP* neuron axons in the antennal lobe labeled with anti-GFP (green) and anti-NCad (magenta). Scale bar, 40 μm. (**C**) Heatmap showing the expression of all detected sensory receptors belonging to the Or (odorant receptor), Gr (gustatory receptor), and Ir (ionotropic receptor) families. A detected receptor is defined as expression at a level of log_2_(CPM+1) ≥ 3 in > 5 cells. (**D**) UMAP plot depicting 26 clusters of adult ORNs. Dimensionality reduction was performed using differentially expressed genes from 42h APF combined with olfactory receptors for a total of 335 genes. (**E–F**) UMAP plots depicting (E) *Or13a* expression in a single cluster (left) and glomerular projection of *Or13a*-positive DC2 ORNs (right), and (F) *Gr21a/Gr63a* expression in a single cluster (left) glomerular projection of *Gr21a/Gr63a*-positive V neurons (right). (**G**) Dot plot depicting sensory receptors used to annotate adult neuron clusters. Clusters containing multiple neuron types are indicated by an underscore separating the names of each type. Each dot represents: mean expression within each cluster (color) and fraction of cells within the cluster expressing the gene (dot size). Asterisk next to VA6_DC3_DM3 and DP1l_DL2d/v_VC3 clusters indicates that some or all of receptors for these cell types are expressed in a small non-overlapping percentage of cells and can be better resolved on a UMAP plot (see Figure 5—supplement 2K, M). (**H**) UMAP plot of 21 clusters annotated to 31 neuron types using antennal chemosensory receptor expression. Note that some clusters contain receptors of multiple ORN types and are labeled as such. The glomerular identity of clusters marked “MARS_#” is unknown. (**I**) Schematic of the glomerular targets of the 31 neuron types mapped in (G–H).

We performed dimensionality reduction using the differentially expressed gene set from 42h APF in combination with antennal olfactory receptors and visualized adult ORN transcriptomes in the UMAP space. ORN transcriptomes were initially divided into 27 clusters; however, we removed a cluster that appeared to contain auditory neurons, leaving us with a total of 26 clusters (Figure 5D; Figure 5—supplement 1A, B). Similar to 24h APF, MARS out-performed density-based clustering of adult ORNs (Figure 5—supplement 1E, F). Though our adult transcriptomes included nuclei labeled with both *unc84-GFP* and *lam-GFP*, neither batch effects nor sequencing depth drove clustering (Figure 5—supplement 1C, D). Finally, the majority of adult cells expressed select odorant binding proteins (Obps) (Figure 5—supplement 1G, H), which are thought to be predominantly expressed in and secreted by accessory cells within a sensillum (Figure 5A’) (Hekmat-Scafe et al., 2002; Larter et al., 2016).

Do adult transcriptomic clusters represent distinct neuron types? We began by evaluating expression of all known antennal sensory receptors. We observed cluster-specific expression of many Ors and Grs in our data (Figure 5E–G, Figure 5—supplement 2). For instance, Or13a (the receptor that DC2-projecting ORNs express) was found in a single cluster (Figure 5E). We could also detect all of the known receptors for V (*Gr21a/Gr63a*), DL3 (*Or65a/b/c*), DC1 (*Or19a/b*), and DM2 (*Or22a/b*) ORN types (Figure 5E, F, Figure 5—supplement 2B, C, J), each of which expresses more than one receptor. Thus, cluster-specific Or and Gr expression allowed us to match 13 transcriptomic clusters with their glomerular types (Figure 5G, H).

In two instances, Ors from multiple ORN types were present in a single cluster. First, we detected expression of *Or49b* (VA5 ORNs) in a distinct set of cells within the larger *Or9a* (VM3 ORNs) cluster (Figure 5G; Figure 5—supplement 2A). Thus, we annotated this cluster as containing both VM3 and VA5 ORN types. Likewise, Ors for DM3 (*Or47a*), VA6 (*Or82a*), and DC3 (*Or83c*) ORNs were found in non-overlapping subsets of cells in a single cluster (Figure 5—supplement 2K). Again, since all three of these Ors were expressed in a distinct portion of cells, we annotated this cluster as representing all three ORN types. We note that in both examples, with the exception of DC3 ORNs, the intermingled ORNs originated from the same sensillar class (Couto et al, 2005). These findings suggest that the transcriptomes of some ORNs within the same sensillar class are indistinguishable at the adult stage, which likely contributed to why we had 26 adult neuron clusters rather than ~50.

We next focused on annotating Ir-expressing clusters. Irs mediate detection of olfactory, humidity, and thermosensory cues (Enjin et al., 2016; Knecht et al., 2017; Budelli et al., 2019). Ir expression is more promiscuous than that of Ors and Grs, as individual Irs can be found in multiple neuron types in the antenna and arista (Grabe et al., 2016; Marin et al., 2020). It was therefore challenging to annotate Ir-positive clusters as belonging to just one neuron type. For instance, *Ir21a*—known to be expressed in neurons projecting from the antenna and arista to the VP1l and VP3 glomeruli, respectively (Marin et al., 2020)—was largely localized to a single cluster (Figure 5G, Figure 5—supplement 2P). Likewise, *Ir40a*—known to be in neurons targeting to the VP4 and VP1d glomeruli (Marin et al., 2020)—was also enriched in one cluster (Figure 5G, Figure 5—supplement 2O). A few cells within the *Ir40a* cluster expressed the arista-specific *Gr28b* (VP2-projecting neurons), indicating that this cluster contained both arista and antennal neurons (Figure 5—supplement 2P). Also, a single cluster expressed *Ir75a/b/c and Or35a* (Figure 5G, Figure 5—supplement 2M), which corresponds to DP1l, DL2d/v, and VC3 ORNs that originate from the same sensillar class (Grabe et al., 2016). Lastly, *Ir75d* (VL1) and *Ir92a* (VM1) ORNs, which are present in the same sensillum (Martin et al., 2013), were found within the same cluster (Figure 5G, Figure 5—supplement 2N). Collectively, we used cluster-specific receptor expression to map 31 antennal sensory neuron types to 20 adult transcriptomic clusters (Figure 5G–I).

### Matching transcriptomic clusters across all stages enabled annotation of additional ORN types

In addition to the 22 annotated ORN types from our prior 42h APF study (Li et al., 2020), we mapped 17 and 31 neuron types to their corresponding 24h APF and adult transcriptomic clusters, respectively (Figure 4G, 5H). When we performed dimensionality reduction using differentially expressed genes from 42h APF and olfactory receptors and visualized neurons of all stages together, we noticed that the transcriptomes readily separated into clusters corresponding to distinct ORN types (Figure 6A, B). The fact that this gene set can accurately differentiate ORNs at all stages indicates that it contains relevant ORN type-specific information. Denoting a high degree of transcriptomic similarity, cells belonging to the same ORN type at 24h and 42h APF predominantly clustered together, whereas adult transcriptomes remained distinct (Figure 6A, B). Differences in either sequencing technology or transcriptome similarity could be the reason adult transcriptomes do not intermingle with pupal ones.

**Figure 6.**
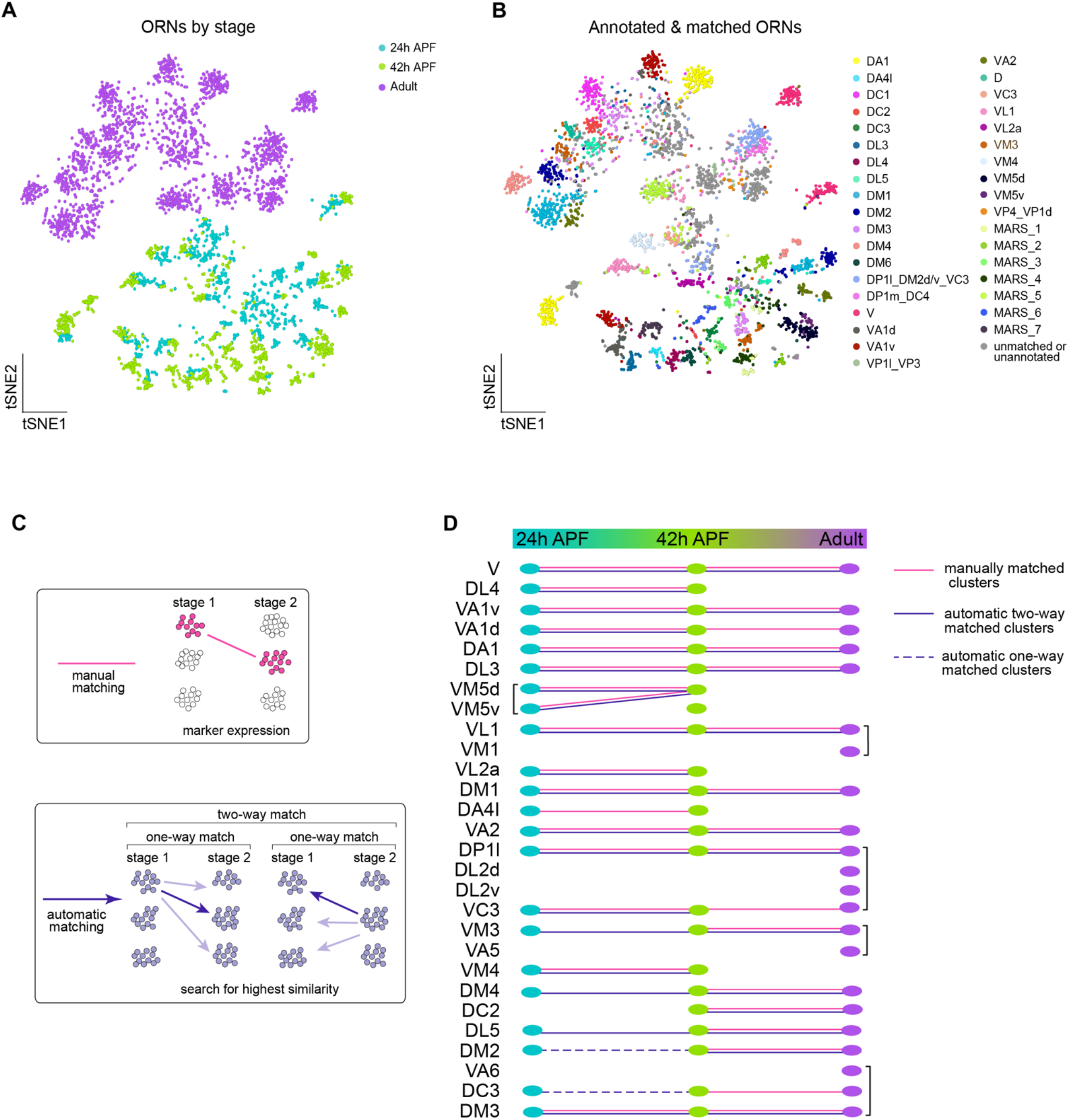
Matching transcriptomic clusters across all stages enabled annotation of additional ORN types. (**A–B**) t-SNE plots of ORNs from the three stages profiled in this study: 24h APF, 42h APF, and adult (A) and annotated and/or matched ORNs across these stages (B). Clusters marked “MARS_#” are matched but their glomerular identity is unknown. Dimensionality reduction was performed using differentially expressed genes from 42h APF combined with olfactory receptors for a total of 335 genes. (**C**) Schematic summarizing the two approaches that were used for matching ORNs from different stages: 1) manual matching using cluster-specific markers/olfactory receptors (top); and 2) automatic matching by calculating the transcriptome similarity between one cluster at a certain stage to all clusters from another stage (bottom). (**D**) Summary of all matched ORN types from 3 different stages, the method used to match clusters is represented by the type of line (color, solid, dashed) linking the clusters between timepoints. Cell types in brackets indicate that they all mapped to a single cluster at that stage.

We next wanted to leverage the similarity between 24h and 42h ORNs, as well as the fact that we have annotated different neuron types between pupal and adult stages, to decode additional clusters at each time point. Since we already employed a manual matching strategy (using marker genes/olfactory receptor expression), we asked if an automatic matching approach that uses transcriptomic similarity to link clusters across development could annotate more clusters (Figure 6C). Thus, we identified differentially expressed genes in each cluster at the 24h APF, 42h APF, and adult stages. Each cluster at one stage was independently compared to every cluster at the other time points. We then computed a similarity score (Jaccard index) for the proportion of genes shared between each pair of clusters across two stages. If two clusters both had the highest similarity score with each other, we considered these clusters to be two-way matched (Figure 6C). One-way matches occurred when one cluster has the highest similarity with a cluster at a different stage, but the opposite did not hold true (Figure 6C, see Methods). Due to the difference in sequencing technologies between pupal and adult ORNs, we did not allow one-way matches between these stages.

We first used this approach to validate our manual matches and found that the results of these two strategies were in complete agreement (Figure 6D). There were certain clusters (e.g., DA4l or VM3) that we were only able to match using one of the strategies (Figure 6D). Can two-way automatic matching allow annotations of additional neuron types that we were unable to link previously? Indeed, this approach matched 6 more clusters from 24h to 42h APF. Two of these clusters corresponded to DM4 and VM3 ORNs and four were unannotated (Figure 6B, D, Figure 6—supplement 1A, C). Furthermore, we annotated the pupal DL5 clusters by matching them to the previously annotated adult DL5 cluster (Figure 6D). For other 42h APF clusters that mapped on to adult clusters containing multiple ORN types, we annotated the adult clusters with the name of the 42h cluster it matched to but noted when there were multiple ORN types within it (Figure 6D, brackets). Lastly, we relied on one-way automatic matching to decode DC3 and DM2 ORNs at 24h APF (Figure 6D) and to link two unannotated types between 24h and 42h APF (Figure 6—supplement 1C). Automatic matching was thus able to decode an additional five transcriptomic clusters at 24h APF and one additional cluster at 42h APF, bringing the total number of annotated clusters at each stage to 21 and 23, respectively. Altogether, by combining manual and automatic approaches, we matched 17 ORN types across all developmental stages.

### ORN transcriptomes in part reflect axon trajectory at 24h APF and sensory modality in adults

Antennal ORNs are located in one of four morphologically distinct sensillar classes: antennal trichoids, antennal coeloconics, large basiconics, and thin/small basiconics (Figure 7A). Thermo- and hygrosensory neurons are found within the sacculus, a three chambered invagination within the antenna, and the arista (Figure 7B) (Enjin et al., 2016; Budelli et al., 2019). Generally, antennal sensory neurons that are found in the same structure (e.g., sensillar class), project their axons to adjacent regions in the antennal lobe, and detect similar stimuli. Since we annotated neurons in each of these sensory structures and matched ORN transcriptomes from each sensillar class across development (Figure 7C, D), we next sought to determine the stage-specific transcriptomic features of neurons in each of these groups.

**Figure 7.**
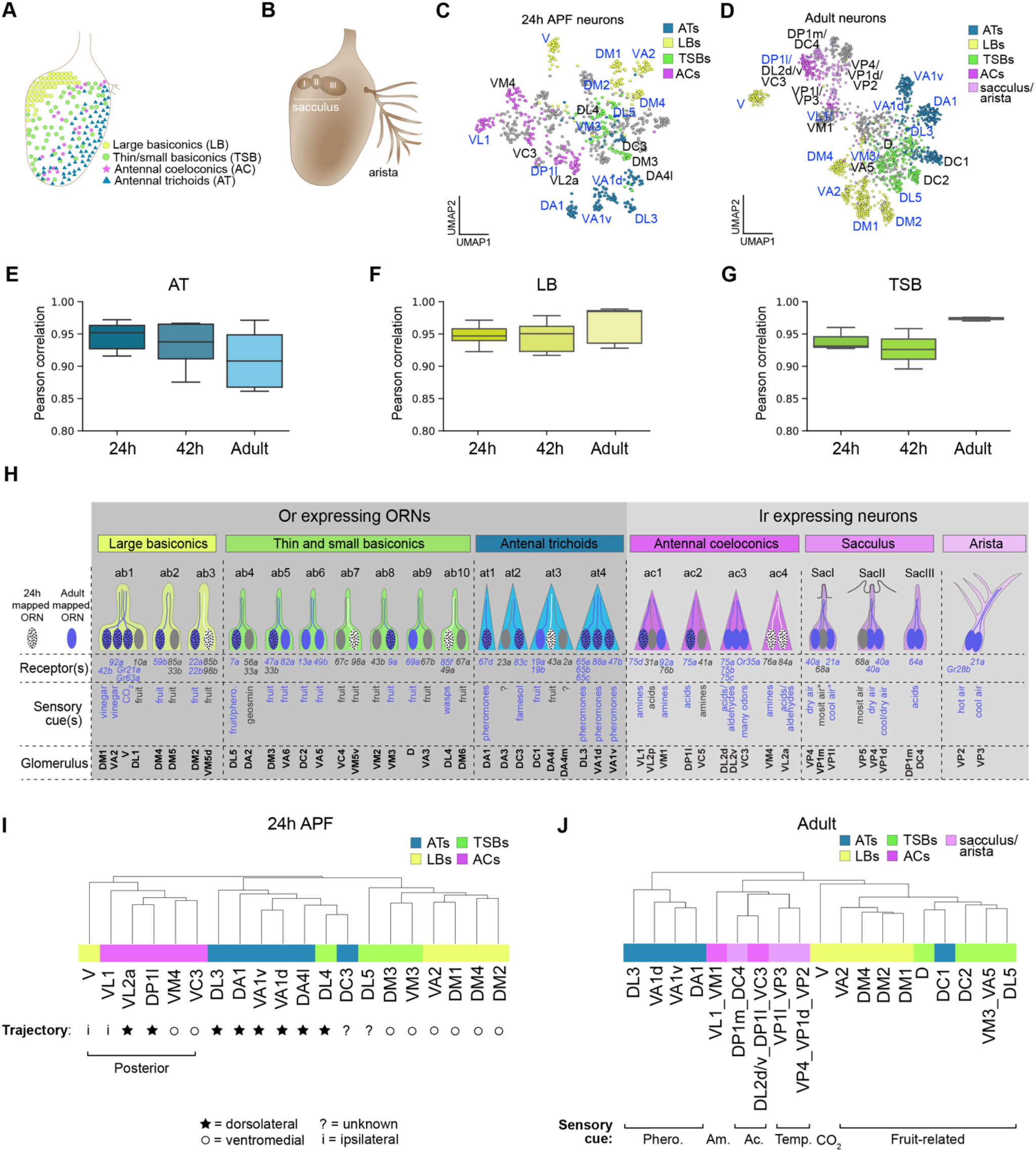
ORN transcriptomes in part reflect axon trajectory at 24h APF and sensory modality in adults. (**A**) Schematic of the antenna depicting the spatial distribution of sensillar classes. (**B**) Schematic of the antenna depicting the structure and location of the sacculus and arista. (**C–D**) UMAP plots of 24h APF (C) and adult neurons (D) colored by sensillar class. Neuron types labeled in blue denote those that are matched at 24h APF, 42h APF, and adult stages. Dimensionality reduction was performed using differentially expressed genes from 42h APF combined with olfactory receptors for a total of 335 genes. (**E–G**) Quantification of cluster-level similarity of neurons in each sensillar class that were matched in Figure 6 at each developmental stage corresponding to: antennal trichoid sensillar class (E), large basiconic sensillar class (F), and thin small basiconic sensillar class (G). 416 DE genes identified from all three stages were used for this analysis. (**H**) Schematic summary of ORNs, thermosensory and hygrosensory neurons within their respective antennal sensillar class/structure, the sensory receptors each neuron type expresses, and the category of cues each type predominantly responds to. White dotted neurons have been mapped to their respective transcriptomic cluster at 24h APF, blue neurons have been mapped to their transcriptomic cluster in adults, and blue dotted neurons have been mapped at all stages. Receptors in blue were detected in either 24h APF or adult datasets. Sensory cues marked with asterisks indicate that they are putative stimuli for this neuron type. See references below for information regarding glomerular projection/receptor expressed (Couto et al., 2005; Fishilevich & Vosshall, 2005; Grabe et al., 2016; Marin et al., 2020) and cues detected (Yao et al., 2005; Hallem & Carlson, 2006; van der Goes van Naters & Carlson, 2007; Kurtovic et al., 2007; Kwon et al., 2007; Semmelhack & Wang, 2009; Silbering et al., 2011; Gallio et al., 2011; Ronderos et al., 2014; Ebrahim et al., 2015; Lin et al., 2015; Enjin et al., 2016; Budelli et al., 2019). (**I–J**) Dendrograms of 24h APF (I) and adult (J) neuron types that resulted from hierarchical clustering of transcriptomes based on differentially expressed genes at each time point. Axon trajectory/projection information is from (Endo et al., 2007; Joo et al., 2013; Silbering et al., 2011; Rybak et al., 2016). The following abbreviations were used to describe the stimuli adult neurons respond to: Phero. = pheromones; Am = amines; Ac = acids; and Temp. = temperature (and humidity sensing VP4 neurons). Some adult clusters contain multiple neuron types, but these clusters are within the same sensillar class or detect the same cues.

We started by measuring cluster-level transcriptomic similarity of matched ORNs within each sensillar class at all stages. (We could not evaluate coeloconic transcriptomes, since we only matched two ORN types within this class.) We found that within each sensillar class, peak cluster-level similarity occurred at different stages (Figure 7E–G). For instance, members of the trichoid class had most the different transcriptomes in adults (Figure 7E). The diversity of signals that different trichoid ORN types respond to, including various pheromones and fruit cues (Kurtovic et al., 2007; Dweck et al., 2013, 2015), could contribute to why these cells have more diverse transcriptomes at this stage. Both basiconic sensillar classes peaked in similarity in the adult stage and were most diverse at 42h APF (Figure 7F, G). In line with this, basiconic clusters contained multiple ORN types at the adult stage (Figure 5H). Removing V neurons, which are distinct in their axon projection pattern and physiological responses (they use Grs as receptors to detect carbon dioxide; Couto et al., 2005; Kwon et al., 2007), did not change the results of the large basiconic comparisons (Figure 7—supplement 1A). With the exception of V neurons, all of the basiconic ORNs we analyzed respond to fruit-related odorants (Semmelhack & Wang, 2009; Dweck et al., 2018), suggesting that transcriptomic similarity is highest between ORNs with similar functions (Figure 7H). Collectively, ORNs within each sensillar class exhibit highest molecular diversity at distinct developmental stages.

Next, we probed the relationship between transcriptome, sensillar class, and stage-related biological processes of neurons at each time point. Previously, we found that 42h APF ORN transcriptomic similarity reflects sensillar class (Li et al., 2020). Thus, we performed hierarchical clustering of all annotated neuron types that mapped to a single sensillar class at 24h APF. Overall, ORNs that belong to the same sensillar class share more similar transcriptomes than those from other classes (Figure 7I). Yet, neither V nor DL4 neurons clustered with their sensillar classes. We therefore considered other factors that could contribute to the clustering of these two ORN types. At 24h APF, most ORN axons are taking either a dorsolateral or ventromedial trajectory around the antennal lobe (Endo et al., 2007; Joo et al., 2013). Could axon trajectory account for the clustering of DL4 and V transcriptomes? Indeed, DL4 ORNs were more similar to the antennal trichoids that take a dorsolateral trajectory than to the ventromedially projecting basiconics annotated here (Figure 7I). Both V neurons and the coeloconics they clustered with project their axons to more posterior glomeruli than the majority of ORNs in trichoid and basiconic classes (Couto et al., 2005; Rybak et al., 2016) (Figure 7I). Posteriorly projecting transcriptomes further segregated based on axon trajectory, as ORNs whose axons take the ventromedial path were more similar to each other than to those taking the dorsolateral trajectory. Further, V and VL1 ORNs have the most divergent transcriptomes of the posteriorly projecting ORNs (Figure 7I), which could reflect the fact that these neurons project their axons ipsilaterally to one antennal lobe and not bilaterally to both lobes, like all the other ORNs in the dendrogram (Couto et al., 2005; Silbering et al., 2011). Moreover, cell surface molecules and transcription factors are important regulators of axon targeting and they constituted 14% and 25% (as opposed to ~6% each in the genome) of the differentially expressed genes used to cluster ORN transcriptomes, respectively (Figure 7—supplement 1B–C). These data suggest that 24h APF ORN transcriptomes reflect similarities in axon trajectory in addition to sensillar class.

In our adult data, we annotated ORNs, thermosensory neurons, and hygrosensory neurons, enabling us to assess the relationship between multiple antennal sensory neuron types at the transcriptome-level. Similar to the pupal stages, neurons within the same sensory structures shared more similar transcriptomes (Figure 7J, Figure 7—supplement 2). Upon closer look, we noticed that adult transcriptomes also reflected the type of stimuli they detect. DC1 ORNs from the trichoid at3 sensillum, for instance, sense limonene (Dweck et al., 2013) and their transcriptomes were more similar to fruit odor-responsive basiconic ORNs than to the pheromone-detecting trichoid ORNs (Figure 7H, J). In fact, DC1 transcriptomes were most similar to D ORNs which can also respond to limonene. Within the Ir-expressing clade, neurons appeared to group by the stimuli they respond to. For example, acid-sensing neurons in the coeloconic sensilla were more closely related to sacculus neurons detecting the same type of stimulus than to amine-sensing coeloconics (Figure 7H, J). Further indicating transcriptomic similarity could reflect sensory modality, the thermo- and hygrosensory neurons in the sacculus and arista were most similar to each other (Figure 7J) and in some cases inseparable at the cluster level (Figure 5H). Taken together, our data suggest that antennal neuron transcriptomic similarities in part reflect stage-related biological processes.

### Transcriptomic differences between Or-expressing and Ir-expressing neurons

Obtaining adult single-cell transcriptomes enabled us to explore gene expression differences between distinct types of antennal sensory neurons. We began by classifying these neurons based on the two major classes of sensory receptors they express: Ors and Irs. Ors are used to detect odorants, whereas Irs can respond to odorants, temperature, and humidity (Hallem & Carlson, 2006; Silbering et al., 2011; Knecht et al., 2016, 2017; Ni et al., 2016). Despite having seven transmembrane domains, insect Ors are not G-protein-coupled receptors (GPCRs) but instead adopt an inverse seven-transmembrane topology with an intracellular amino-terminal domain and an extracellular carboxy-terminal domain (Figure 8A) (Benton et al., 2006). Ors assemble into heterotetramers with their obligate co-receptor, Orco, to form non-selective, ligand-gated cation channels (Butterwick et al., 2018). In our adult transcriptomes, 69% of neurons express *Orco* (Figure 8B, Figure 8—supplement 1A). Irs are glutamate insensitive relatives of ionotropic glutamate receptors (iGluRs) and are proposed to exist as heterotetrameric cation channels with their co-receptors (Figure 8A) (Abuin et al., 2019). Expression of the *Ir8a* and *Ir76b* co-receptors was restricted to Ir-positive clusters, whereas the *Ir25a* and *Ir93a* co-receptors were expressed in both Or- and Ir-positive neurons in our data (Figure 8C).

**Figure 8.**
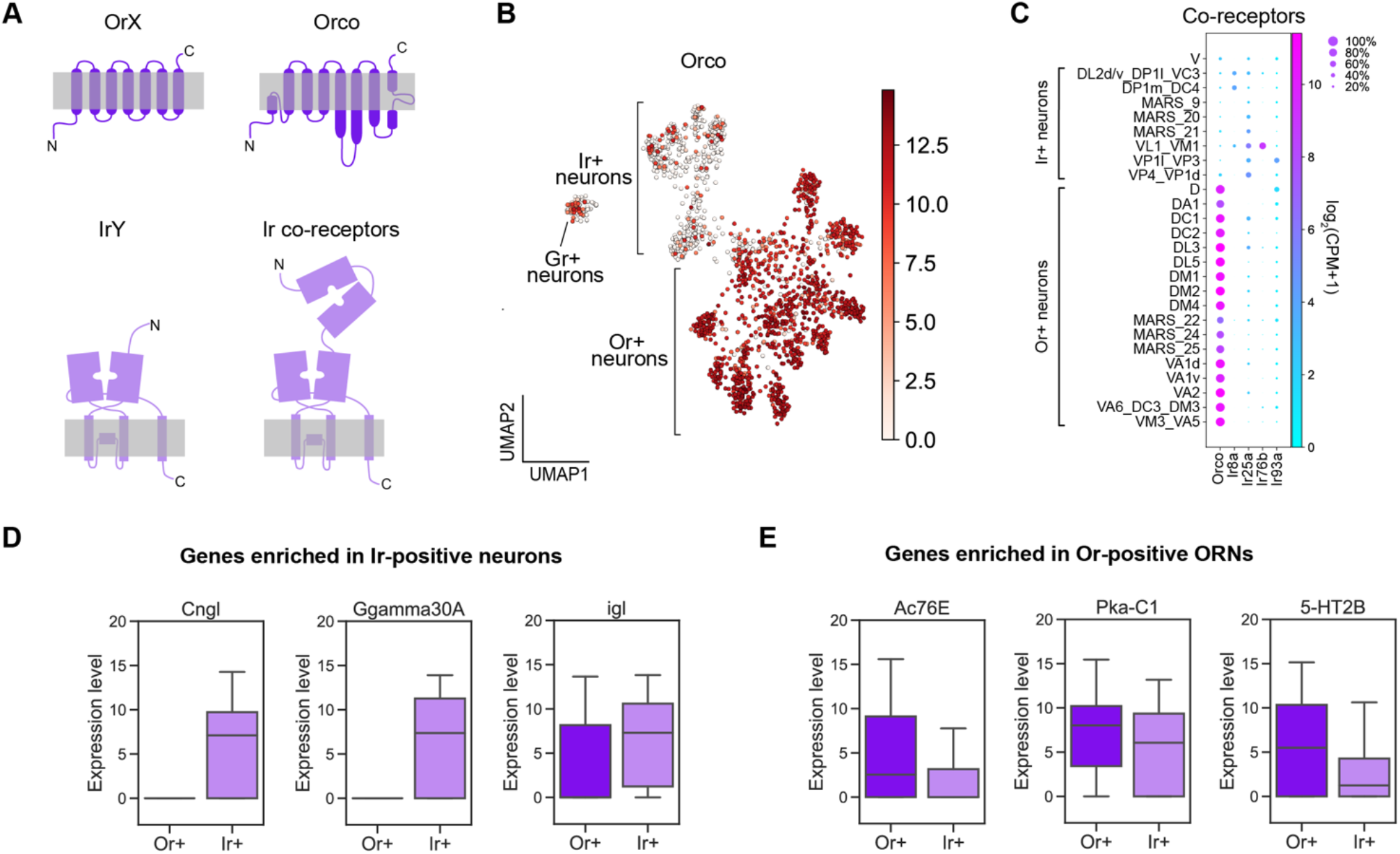
Transcriptomic differences between Or-expressing and Ir-expressing antennal neurons. (**A**) Schematic of two classes of *Drosophila* sensory receptors. OrX (odorant receptors) and their co-receptor Orco are seven-pass transmembrane receptors (top). IrY (ionotropic receptors) and their co-receptors are relatives of ionotropic glutamate receptors. Orco diagram is based on Butterwick et al., 2018 and Ir/Ir co-receptor diagram is based on Abuin et al., 2019. (**B**) UMAP plot depicting *Orco* expression in adult antennal neurons. Dimensionality reduction was performed using previously described methods. Ir-positive and Or-positive classifications were based on both sensory receptor expression and *Orco* expression level within a cluster. Since *Orco* is known to be a Or co-receptor we used high *Orco* expression reveal Or-positive clusters that did not express an identifying receptor, and *Orco* low clusters were annotated as Ir-positive. (**C**) Dot plot depicting Or and Ir co-receptor expression in adult clusters. Clusters containing multiple neuron types are noted using an underscore to separate names of each type. MARS clusters were included because *Orco* expression within these clusters indicates whether they should express an Or or Ir. Each dot represents: mean expression within each cluster (color) and fraction of cells within the cluster expressing the gene (dot size). (**D–E**) Box plots depicting differentially expressed genes between Ir-positive and Or-positive clusters.

Aside from the receptors and co-receptors, little is known about signaling mechanisms that distinguish Or- and Ir-expressing neurons. To examine this, we identified 262 differentially expressed genes (p < 10^−5^) between Or- and Ir-positive neurons (Figure 8—supplement 1B). Although both Irs and Ors are cation channels, we observed differential expression of genes associated with GPCR signaling cascades between these two groups. For instance, Ir-positive neurons preferentially expressed a specific gamma subunit of the heterotrimeric G-protein complex *Ggamma30A*, the calmodulin binding protein *igl*, and the cyclic nucleotide-gated channel subunit *Cngl* (Figure 8D). The adenylyl cyclase *Ac76E*, which belongs to a class of enzymes that often acts downstream of GPCRs, was enriched in Or-positive ORNs (Figure 8E). Additionally, the catalytic subunit of the cAMP-dependent protein kinase, *PKA-C1*, was expressed in both Or- and Ir-positive neurons (Figure 8E). The function of G-protein, cyclic nucleotide, and calmodulin signaling in ORNs is unclear, as some studies find that Or-positive ORNs utilize them for signaling (Wicher et al., 2008) but others report that they do not (Kain et al., 2008; Yao & Carlson, 2010). Additionally, whether Irs engage these signaling mechanisms is unknown. Our data suggest that both Or- and Ir-expressing neurons may utilize distinct components of GPCR signaling.

GPCR signaling could be employed by neuromodulators or neurotransmitters that use GPCRs as receptors. ORNs not only synapse with PN dendrites, but are also contacted by interneurons of various neurotransmitter profiles whose signaling could engage distinct GPCRs on ORN terminals (Olsen et al., 2007; Roy et al., 2007; Shang et al., 2007; Chou et al., 2010). We found that the serotonin receptor *5-HT2B* had higher expression in Or-positive clusters compared to Ir-positive clusters (Figure 8E). By contrast, *5-HT1A* was expressed in a subset of Ir-positive clusters (Figure 8—supplement 1B). Given that 5-HT2B promotes excitability whereas 5-HT1A reduces it (Tierney, 2018), these data suggest that serotonin could have differential effects on Ir- and Or-expressing neurons.

### Differential expression of ion channels, neurotransmitter receptors, and neuropeptides among distinct sensory neuron types

Finally, we investigated the expression of genes that regulate neuronal excitability and synaptic transmission among different sensory neuron types. First, we queried expression of neuropeptides. Short neuropeptide F precursor (*sNPF*), which regulates olfactory sensitivity of DM1 ORNs (Root et al., 2011), was found in all adult clusters but was more highly expressed by Ir-positive neurons (Figure 9A). Ion transport peptide (*ITP*) displayed expression in a few Or-positive clusters (Figure 9A). Next, we assayed expression of GPCRs that serve as neurotransmitter, hormone, and neuropeptide receptors. Many of these receptors had broad expression in multiple neuron types, such as the dopamine receptor (*Dop1R1*), octopamine receptors (*Octβ3R, Octβ1R*), neuropeptide receptor (*TrissinR*), metabotropic GABA receptor (*GABA-B-R1*), dopamine/ecdysteroid receptor (*DopEcR*), and muscarinic acetylcholine receptor (*mAChR-B*) (Figure 9A). Expression of other GPCRs was restricted to a subset of clusters, such as the short neuropeptide F receptor (*sNPF-R*), octopamine receptor (*Octβ2R*), allatostatin C receptor 2 (*AstC-R2*), and sex peptide receptor (*SPR) (*Figure 9A). Furthermore, the diuretic hormone receptors, *Dh44-R1* and *Dh31-R*, were mainly expressed by pheromone-detecting DA1 and DA1/DL3 ORNs, respectively (Figure 9A). These data indicate that ORNs express a breadth of GPCRs and may be modulated by a variety of neurotransmitters and hormones.

**Figure 9.**
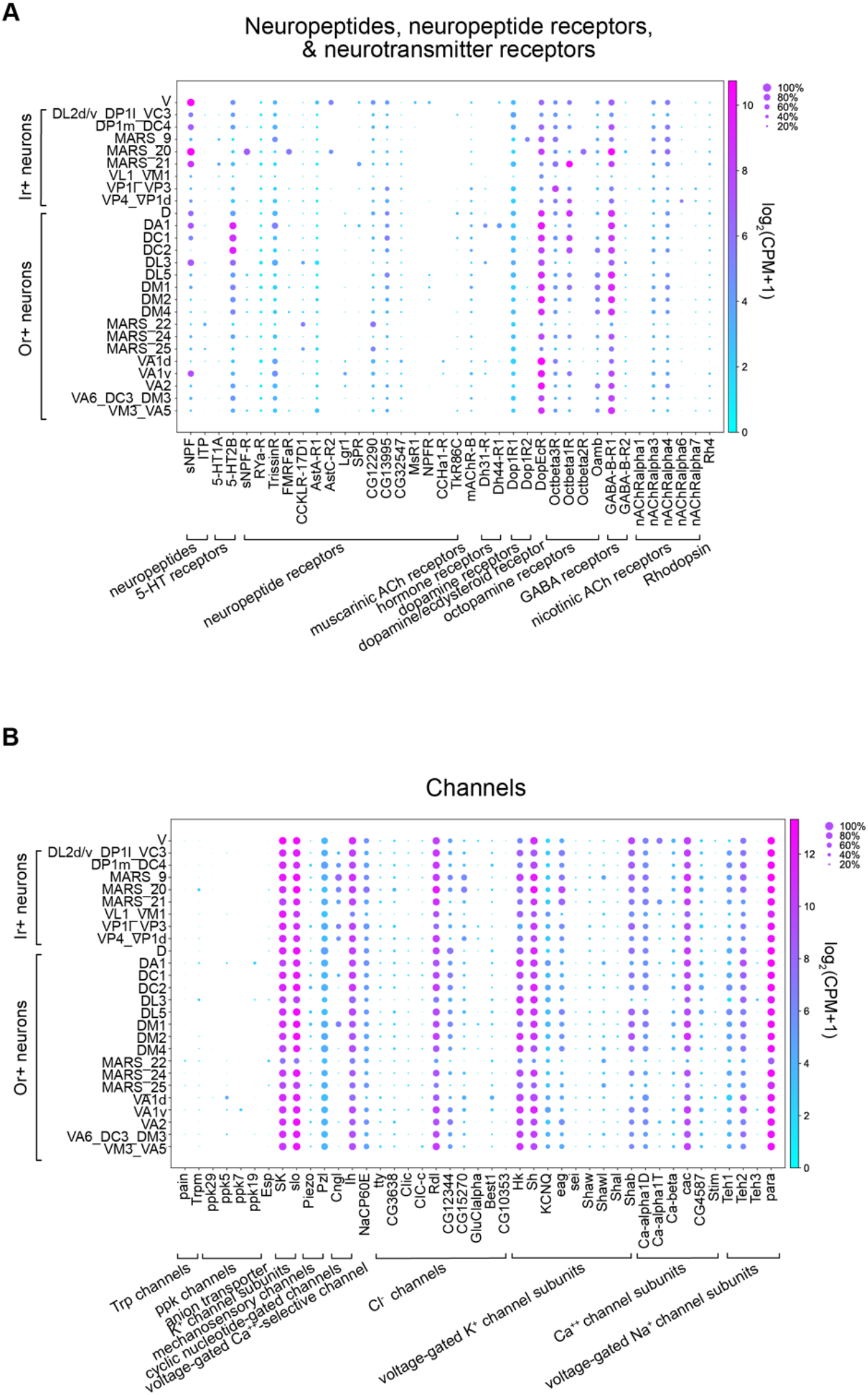
Differential expression of ion channels, neurotransmitter receptors, and neuropeptides among distinct sensory neuron types. (**A–B**) Dot plots depicting expression of neuropeptides, neuropeptide receptors, neurotransmitter receptors, and hormone receptors (A) and ion channels and channel subunits (B) in adult clusters. Clusters containing multiple neuron types are noted using an underscore to separate names of each type. MARS clusters were included because *Orco* expression indicates whether the cluster should express an Or or Ir. Each dot represents: mean expression within each cluster (color) and fraction of cells within the cluster expressing the gene (dot size).

Next, we determined the expression of additional groups of genes important for neuronal physiology. We began with ionotropic neurotransmitter receptors and found that two nicotinic acetylcholine receptor subunits, *nAChRα3* and *nAChRr4*, were broadly expressed in all clusters (Figure 9A). Conversely, *nAChRα1, nAChRα6*, and *nAChRα7* exhibited lower albeit cluster-specific expression in some Ir-positive cell-types (Figure 9A), indicating that distinct subunits could comprise nicotinic acetylcholine receptors in these neurons. Finally, we evaluated ion channel and channel subunit expression in adult transcriptomic clusters. Both pickpocket (ppk) and transient receptor potential (Trp) channel members showed low but restricted expression to a few, mostly trichoid clusters (VA1v, VA1d, DL3; Figure 9B). The rest of the channels and their accessory subunits exhibited broad expression, with the exception of a β subunit of a voltage-gated sodium channel (*Teh3*), an *α* subunit of calcium channel (*Ca-α1T*), a glutamate-gated chloride channel *α* subunit (*GluClα*), and the cyclic nucleotide gated ion channel (*Cngl*) (Figure 8D, 9B). Taken together, these analyses exemplify how our adult transcriptomic data can be used as a foundation to investigate physiological functions of these sensory neurons.

## Discussion

Here, we generated high-quality transcriptomes of ~3,100 sensory neurons from the *Drosophila* third antennal segment at 24h APF and in adults. We integrated these data with the transcriptomes of mid-pupal ORNs (Li et al., 2020). Our clustering methods separated 24h APF and adult transcriptomes into 36 and 26 clusters, respectively, and further divided 42h APF transcriptomes into 34 clusters. Importantly, we matched 16 annotated ORN types at all three developmental stages. The fact that we observed fewer clusters than the ~50 glomerular types could be the result of: 1) not enough cells being profiled for some types; 2) closely related types forming a single cluster (e.g., VM5d and VM5v ORNs at 24h APF or VA5 and VM3 ORNs in adults; Figure 4G, 5H); or 3) some Ir-positive neuronal types expressing the same receptor and cannot be further subdivided (Figure 5H; Figure 5—supplement 2M, N, P). Our analysis of sensory receptor and marker expression within clusters points to the latter two explanations accounting for the majority of the reduction in clusters compared to the number of neuron types. We were also unable to annotate thermo- and hygrosensory neurons in our pupal data, possibly because these cells both exist in low numbers in the antenna and did not express distinguishing receptors in pupal transcriptomes. Nonetheless, we annotated the majority of clusters at all three stages and found that they represented anatomically and functionally distinct sensory neuron types.

### Single-nucleus RNA-seq can be used to profile cuticle-associated tissues

Single-cell transcriptomics is a powerful tool for the discovery of novel cell types, revealing subtype specific markers, and studying gene expression changes underlying cellular development and maturation (Li, 2020). Adult ORNs, like many insect tissues, are tightly associated with the hardened exoskeleton and are difficult to isolate using standard cell dissociation protocols. Nuclei, on the other hand, are much more resistant to mechanical assaults and have been readily profiled using snRNA-seq in multiple systems (Grindberg et al., 2013; Habib et al., 2016, 2017; Cui et al., 2020; Kebschull et al., 2020; Lau et al., 2020). snRNA-seq has many advantages including: 1) tissue can be kept in long-term storage at −80°C prior to extraction (Wu et al., 2019; Ding et al., 2020); 2) snRNAseq reduces cell-type sampling bias (Denisenko et al., 2020); and 3) nuclei are more easily captured by microfluidics devices than large and/or fragile cells (Cui et al., 2020; Denisenko et al., 2020). Underscoring the utility of this protocol, high-quality nuclei can be isolated from frozen tissues which allows researchers studying low-abundance cell types to collect and freeze a large amount of tissue (over multiple days) to ensure that they have an adequate number of nuclei for downstream processing and analyses.

Here, we directly compared snRNA-seq with scRNA-seq of both peripheral and central neurons: 24h APF ORNs and adult PNs, respectively. Similar to a previous report in mammalian cortical neurons (Bakken et al., 2018), we observed a reduction in genes detected in nuclei compared to stage-matched cells. Including intronic reads in our nuclear transcriptomes increased gene detection rates in both neuron types (Figure 2—supplement 1 D, E). Critically, we found that our scRNA-seq and snRNA-seq protocols detected largely overlapping genes (Figure 2K, L). Finally, we showed that this method could be used to separate closely-related adult transcriptomes (Figure 5). Collectively, we demonstrated that snRNA-seq is well-suited for profiling previously inaccessible neurons in the adult fly and this protocol can be easily used by other researchers that want to gain transcriptomic access to their tissue of interest (see protocol in Methods section).

### ORN transcriptomic similarity reflects multiple biological factors

Understanding the molecular underpinnings of neural circuit formation, maintenance, and function requires that we can access the transcriptomes of anatomically and functionally distinct neuronal types at developmentally-relevant stages. Here, we created a dataset that will enable these types of analyses by profiling *Drosophila* ORNs at stages corresponding to: 1) initial axon pathfinding (24h APF); 2) synaptic partner matching (42h APF; Li et al., 2020); and 3) sensory function (adult).

Cluster-level comparison within each newly profiled stage lead to the discovery that type-specific transcriptomes reflect multiple biological factors. Initially, the majority of 24h APF ORNs appeared to reflect their sensillar class; however, neither DL4 nor V neurons fit this pattern (Figure 7I). Rather, transcriptomes of both DL4 and V projecting neurons were most closely related to ORNs whose axons take similar trajectories. Future work decoding additional ORN types could enable a more complete evaluation of the degree to which 24h APF transcriptomes reflect axon trajectory and sensillar class.

In our adult data, we annotated olfactory, thermosensory, and hygrosensory neurons originating from all of the sensory structures of the antenna. Neurons housed within the same structure generally detect similar stimuli, with a few exceptions (Figure 7H). Thus, when we compared adult transcriptomes we found that most neurons housed in the same sensory structure were very similar to each other. However, when we queried the exceptions to this clustering (e.g., DC1 and some coeloconic ORNs) adult transcriptomes appeared to also reflect similarities in the sensory stimuli they detect (Figure 7J). As an example, coeloconic neurons that detect acids are more similar to acid detecting saccular neurons than to coeloconics that sense amines (Figure 7J). While it is not possible to cleanly separate the stimulus a neuron type detects from the sensory structure it is housed in, our data suggests that neurons that detect similar cues may require analogous genes for their function. Together, our 24h APF and adult analyses revealed that stage-specific transcriptomes reflect multiple factors and underscore that profiling cells at multiple stages can deepen our understanding of their biology.

### Transcriptomic differences could reveal regulators of adult neuronal function

ORNs employ two main receptor types for the detection of sensory stimuli: Ors and Irs. Both receptor types are thought to form ion channels with their co-receptors (Butterwick et al., 2018; Abuin et al., 2019). However, a metabotropic component to Or signaling has been reported (Wicher et al., 2008). In fact, it has been proposed that Ors are metabotropically-modulated ionotropic receptors; however, the mechanism by which this occurs is unresolved (Nakagawa & Vosshall, 2009; Gomez-Diaz et al., 2018). We found that Or- and Ir-positive neurons express distinct components of GPCR and downstream signaling cascades. Whether these components are used to modulate Or (or Ir) function remains to be determined.

GPCR-directed signaling mechanisms could also be activated by receptors for neuromodulators, neuropeptides, or neurotransmitters. We showed that ORNs express a rich repertoire of neuropeptide, amine, hormone, and metabotropic GABA receptors, some of which are differentially expressed in distinct neuron types (Figure 9A). What is the source of the ligands for these GPCRs? Morphologically and functionally diverse local interneurons send processes to the antennal lobe where they contact ORN axons and PN dendrites (Chou et al., 2010). While most local interneurons are GABAergic, some are cholinergic or glutamatergic, and a few are dopaminergic (Olsen et al., 2007; Shang et al., 2007; Chou et al., 2010). Local interneurons could also signal to ORNs via secreted hormones or neuropeptides. Furthermore, biogenic amines are known to modulate ORNs in *Drosophila* and other insect species (Dolzer et al., 2001), and deuterocerebral neurons and ventral unpaired neurons providing serotoninergic and octopaminergic inputs into the antennal lobe, respectively (Roy et al., 2007; Sinakevitch & Strausfeld, 2006; Zhang & Gaudry, 2016). It has been reported that serotonin from deuterocerebral neurons does not directly affect ORNs (Dacks et al., 2009), but whether these cells modulate Ir-positive neurons is not known. In other insects, both serotonin and octopamine modulate odor-evoked responses and are found in hemolymph, the fluid bathing all organs including the antenna (Dolzer et al., 2001; Grosmaitre et al., 2001; French et al., 2014; Zhang & Gaudry, 2016). Thus, identifying the subcellular localization of these aminergic receptors will provide important insight into their function in ORNs.

In conclusion, by profiling *Drosophila* ORNs at multiple stages, we have generated a rich dataset that can be used as a foundation to reveal molecular regulators of neuron differentiation, axon targeting, synapse formation, and sensory function. This work, together our accompanying paper profiling PNs at multiple stages (Xie et al.), should spur novel and important biological discoveries.

## Materials and Methods

### Key Resources Table

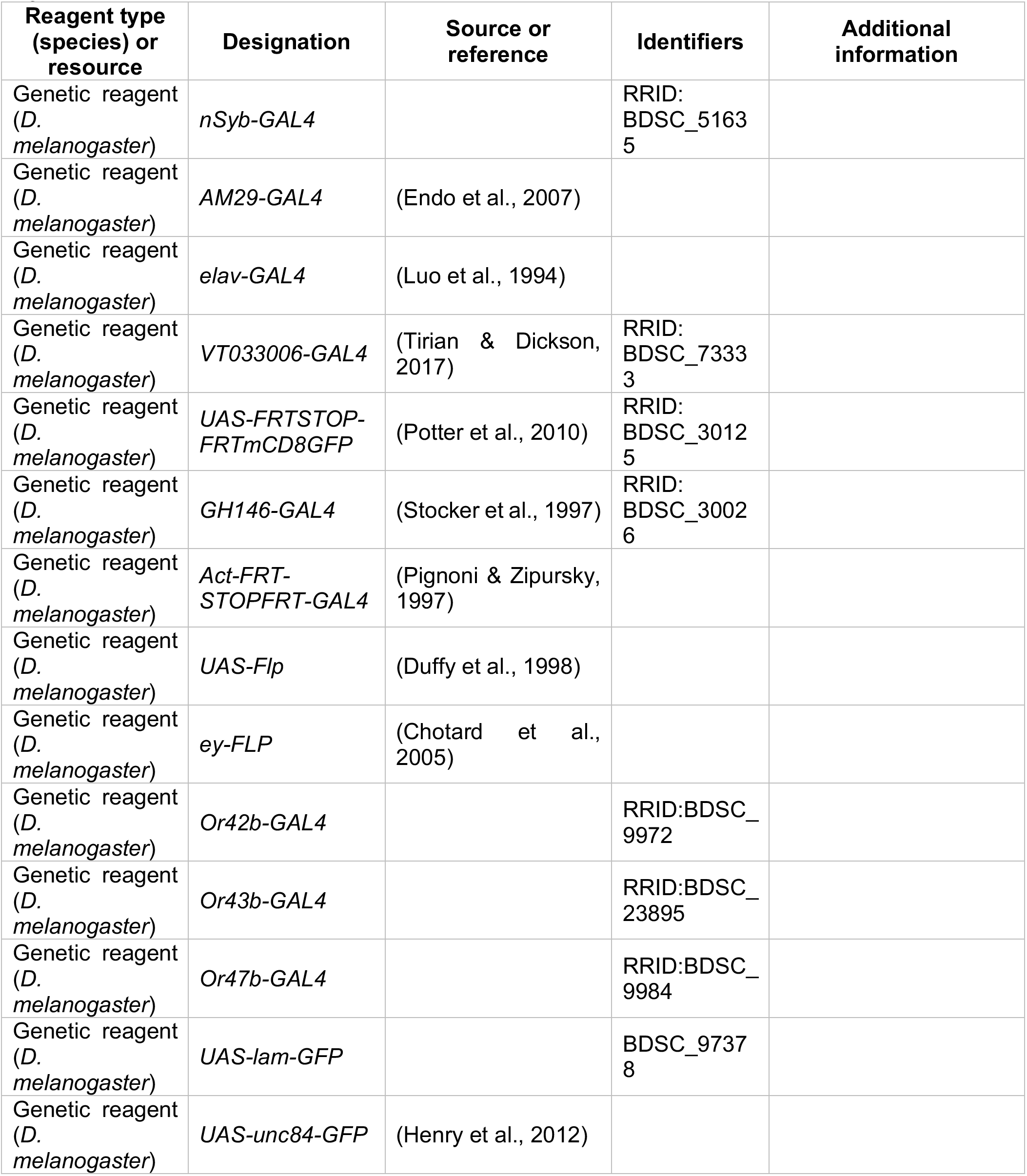
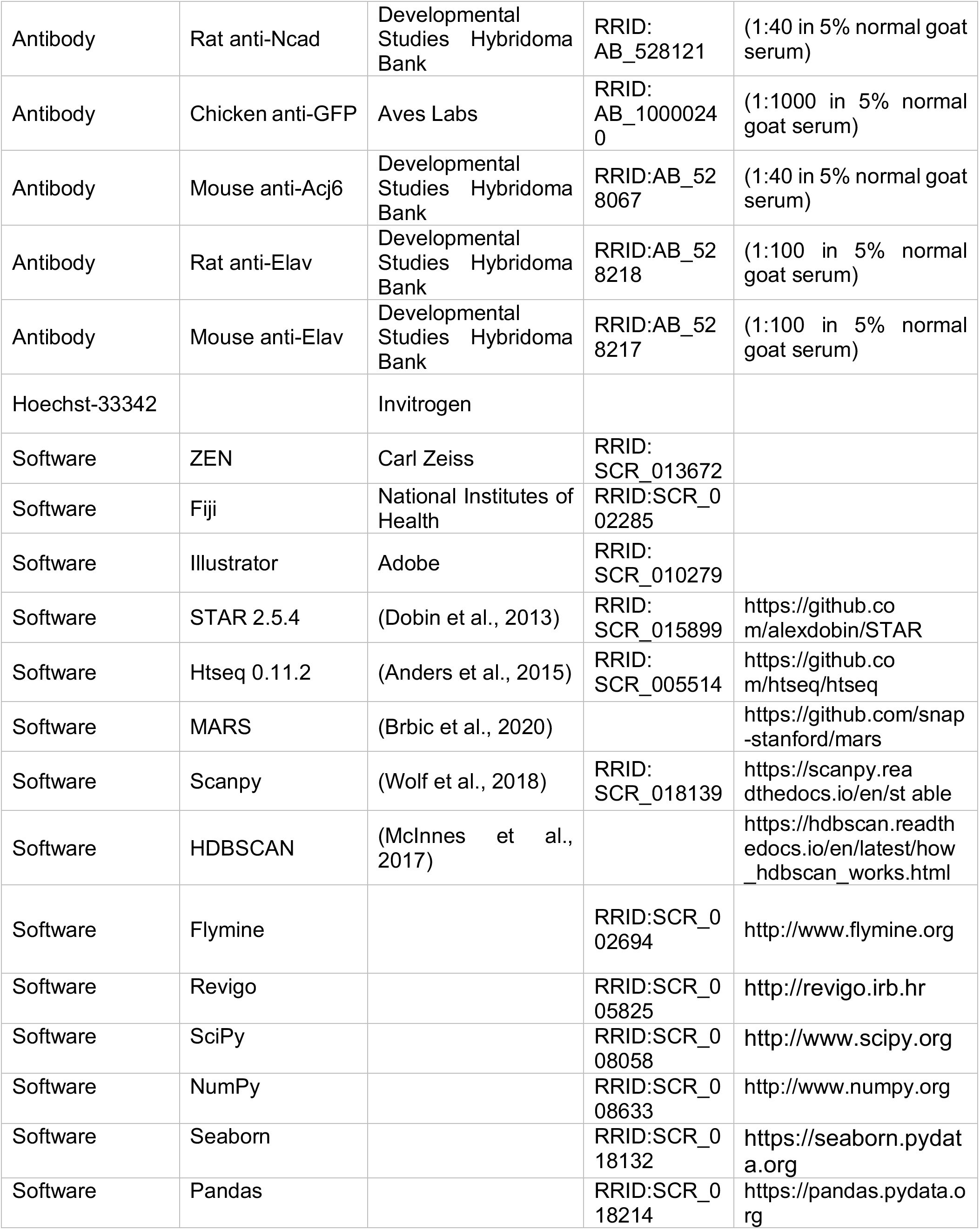
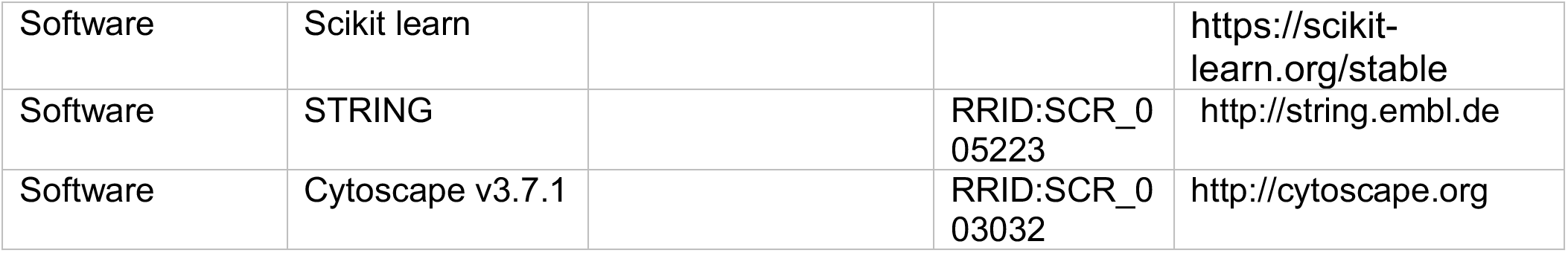

### *Drosophila* stocks and genotypes

Flies were maintained at 25°C on standard cornmeal medium with at 12-hour light-dark cycle. The following fly lines were used in this study: *nSyb-GAL4* (Bloomington *Drosophila* Stock Center, BDSC #51635); *AM29-GAL4* (Endo et al., 2007); *elav-GAL4* (Luo et al., 1994); *UAS-FRT-STOP-FRT-mCD8-GFP* (Potter et al, 2010)*; ey-FLP* (Chotard et al., 2005); *VT033006-GAL4* (Tirian & Dickson, 2017); *Or42b-GAL4* (BDSC #9972); *Or43b-GAL4* (BDSC #23895), *Or47b-GAL4* (BDSC #9984); *GH146-GAL4* (Stocker et al., 1997); *UAS-lam-GFP* (BDSC #97378); *UAS-unc84-GFP* (Henry et al., 2012); UAS-Flp (Duffy et al., 1998); *Act-FRT-STOP-FRT-GAL4* (Pignoni & Zipursky, 1997). Note: 48h APF and adult cells/nuclei were labeled with *nSyb-GAL4*; however, this driver was not sufficiently expressed at 24h APF so cells and nuclei from this earlier stage were labeled using *elav-GAL4*. Also, PN cells were labeled with *GH146-GAL4* and nuclei with *VT03006-GAL4*. For the permanent labeling experiments, *Or-GAL4* lines were crossed with permanent labeling ready flies, *UAS-Flp, Act-FRT-STOP-FRT-GAL4, UAS-CD8-GFP*, and F1 adult brains were dissected and analyzed.

### Immunofluorescence and image acquisition

Fly brains and antennae were dissected and labeled according to previously described methods (Wu & Luo, 2006; Li et al., 2020). Briefly, brains and antennae were rapidly dissected in PBS and put into ice cold 4% paraformaldehyde in PBS, washed and permeabilized in PBST (1X PBS, 0.3% Triton-X-100), incubated in blocking buffer (5% normal goat serum in PBST), incubated in primary antibody overnight at 4°C, washed in PBST, and incubated in secondary antibody overnight at 4°C. The following primary antibodies were used in this study: rat anti-Ncad [N-Ex #8; at 1:40; Developmental Studies Hybridoma Bank (DSHB)]; chicken anti-GFP (1:1000; Aves Labs); mouse anti-Elav (9F8A9; 1:100; DSHB); rat anti-Elav (7E8A10; 1:100; DSHB); mouse anti-Acj6 (1:40; DSHB). Secondary antibodies raised in goat or donkey against mouse, rat, or chicken antisera and conjugated to Alexa Fluor 488/594/647 (Jackson ImmunoResearch) were used at 1:200. Tissue was mounted in SlowFade Gold (Thermo Fisher Scientific). 16-bit images were acquired using a 20x Plan-apochromat (0.8 NA) or 40x Flaur (1.3 NA) objective on a Zeiss LSM 780 (Carl Zeiss) using Zen software (Carl Zeiss) and processed with Fiji (ImageJ; National Institutes of Health)

### Single-cell RNA-seq

Single-cell RNA-seq *of mCD8-GFP* labeled ORNs was performed following the protocol we previously described (Li et al., 2017). Briefly, *Drosophila* third antennal segments labeled with mCD8-GFP using specific GAL4 drivers were manually dissected. Single-cell suspensions were then prepared. Individually labeled cells were sorted via fluorescence-activated cell sorting (FACS) into single wells of 384-well plates containing lysis buffer using an SH800 instrument (Sony Biotechnology). Full-length poly(A)-tailed RNA was reverse-transcribed and amplified by PCR following the SMART-seq2 protocol. To increase cDNA yield and detection efficiency, we increased the number of PCR cycles to 25. Primer dimer artifacts were reduced by digesting the reverse-transcribed first-strand cDNA using lambda exonuclease (New England Biolabs) (37°C for 30 min) prior to PCR amplification. Sequencing libraries were prepared from amplified cDNA using tagmentation (Nextera XT). Sequencing was performed using the Novaseq 6000 Sequencing system (Illumina) with 100 paired-end reads and 2 x 12 bp index reads.

### Single-nucleus RNA-seq

Single-nucleus RNA-seq was performed using the same SMART-seq2 protocol as single-cell RNA-seq with a few key modifications. First, nuclei are labeled with a fluorescent nuclear marker instead of a membrane-bound one. Second, the dissociation and single-nucleus suspension protocol varied greatly from that used for single-cells, detailed below, and was adapted from Krishnaswami et al., 2016. Third, cDNA was amplified with 28 PCR cycles (instead of 25) and reads were aligned to exons and introns.

Single-nucleus suspensions were made using the following protocol:

1. Cross desired GAL4 lines with UAS-nuclear-GFP. Do not use *UAS-nlsGFP*, because it does not give a GFP signal after nucleus isolation. We instead use *UAS-unc84-GFP* and *UAS-lam-GFP*.
2. Dissect tissue in cold Schneider’s medium, and use P20 pipet (prior to beginning, coat the tip with extraneous tissues) to transfer tissue into 100 μl Schneider’s medium in a nuclease-free 1.5ml tube on ice. Label the tube clearly using permanent marker. **Note**: for tissues that float in the medium (e.g., adult antennae), before dissection, prepare three clean dishes: first with 100% ethanol, second and third with Schneider’s medium. Rinse the fly in the first dish with 100% ethanol for 5 seconds, then rinse the fly in the second dish, and dissect in the third dish.
3. After dissection, spin samples down using bench top centrifuge.
4. **Fresh**: The sample can be processed for nuclei extraction immediately following dissection.
5. **Frozen**: Alternatively, the sample can be flash-frozen for long-term storage. Seal the 1.5 ml EP tube with parafilm and put into liquid nitrogen for >30s. Immediately store the sample in −80°C freezer for long-term storage (several months).
6. Prepare fresh homogenization buffer and keep on ice using the following recipe:
7. Thaw samples from −80°C on ice if using frozen samples. Centrifuge samples in 100 μl Schneider’s medium using bench top centrifuge, discard medium, and add 100 μl homogenization buffer.
8. Optional: if sample pieces are too big, e.g., whole body/head, use pellet pestle motor (Kimble 6HAZ6) to grind the sample for 30s–60s on ice.
9. Add 900 μl homogenization buffer, and transfer 1000 μl homogenized sample into the 1 ml dounce (Wheaton #357538). Dounce set should be autoclaved at 200°C >5h.
10. Release nuclei by 20 strokes of loose Dounce pestle and 40 of tight Dounce pestle. Keep on ice. Avoid bubbles.
11. Filter 1000 μl sample through 5 ml cell strainer (35 μm), and then filter sample using 40 μm Flowmi (BelArt #H13680-0040) into 1.5 ml EP tube.
12. Centrifuge for 10 min at 1000 g at 4°C. Discard the supernatant. Do not disturb the pellet.
13. Re-suspend the nuclei using desired amount (we normally use 500–1000 μl) of 1xPBS/0.5%BSA with RNase inhibitor (9.5 ml 1x PBS, 0.5 ml 10% BSA, 50 μl RNasin Plus). Pipet > 20 times to completely re-suspend the nuclei. Filter sample using 40 μm Flowmi into a new 5 ml FACS tube and keep the tube on ice.
14. Nuclei are stickier than whole cells. For users making single-nucleus suspension for the first time, we suggest taking 10 μl of the single-nucleus suspension, stain with Hoechst-33342 (Invitrogen), and check on a cell counter slide to confirm if they are mostly individual nuclei. If nuclei are not sufficiently dissociated, adjust above steps (e.g., increase the number of strokes of the tight pestle when releasing nuclei).
15. Collect nuclei using FACS. Set up the gates using 4 groups of nuclei: WT nuclei, GFP nuclei, WT nuclei + Hoechst stain, GFP nuclei + Hoechst stain. For different fly tissues, there may be more than one band of nuclei with varying Hoechst intensities. They represent different cell types with different nuclei sizes (polyploidy is common for many fly tissues). It is advised to collect different bands of nuclei and check them under microscope. Collect single nuclei into 384-well plates for Smart-seq2, or into a tube for 10X Genomics (not used in this study; details can be sent upon request).

|  | Amount | Storage | Item (add in this order) | Final concentration |
| --- | --- | --- | --- | --- |
| 1 | 10 ml | RT | H<sub>2</sub>O (nuclease free) |  |
| 2 | 0.856 g | RT | Sucrose | 250 mM |
| 3 | 100 μl | 4$^{\circ}\text{C}$ | 1M Tris pH 8.0 | 10 μM |
| 4 | 250 μl | 4$^{\circ}\text{C}$ | 1M KCl | 25 mM |
| 5 | 50 μl | 4$^{\circ}\text{C}$ | 1M MgCl<sub>2</sub> | 5 mM |
| 6 | 100 μl | 4$^{\circ}\text{C}$ | 10% Triton-X 100 | 0.1% |
| 7 | 50 μl | $-20^{\circ}\text{C}$ | RNasin Plus (Promega, N2615) | 0.5% |
| 8 | 200 μl | $-20^{\circ}\text{C}$ aliquots | 50x protease inhibitor (Promega, G6521) | 1x |
| 9 | 50 μl | $-20^{\circ}\text{C}$ | 20mM DTT | 0.1 mM |

### Sequence alignment and preprocessing

Reads were aligned to the *Drosophila melanogaster* genome (r6.10) using STAR (2.5.4) (Dobin et al., 2013). Gene counts were produced using HTseq (0.11.2) with default settings except “-m intersection-strict” (Anders et al., 2015). Gene counts of scRNA-seq were generated using exonic GTF files. Gene counts of snRNA-seq were generated using both exonic and intronic GTF files, and the two gene count tables were merged for analysis. Low quality cells/nuclei having fewer than 50,000 uniquely mapped reads were removed. Sequencing depth was normalized across individual cells by re-scaling gene counts to counts per million reads (CPM). All analyses were performed after converting gene counts to log_2_(CPM+1). Non-neuronal cells were filtered out by selecting for the cells with expression of at least 2 of 5 neuronal markers (*elav, brp, CadN, nSyb, Syt1*) at log_2_(CPM+1) ≥ 2.

### Dimensionality reduction and HDBSCAN clustering

Genes for dimensionality reduction were selected using previously described over-dispersion methods (Li et al., 2020) or by calculating differentially expressed genes between all of the time points (Figure 1F), or within a single time point (Figures 4G, 5H, 6A,B). Note the differentially expressed genes that were used for dimensionality reduction of each individual time point (Figure 4–5) and all time points in Figure 6 were identified at 42h APF and combined with known antennal olfactory receptors from (Grabe et al., 2016) for a total of 335 genes. Principal component analysis (PCA) followed by t-distributed Stochastic Neighbor Embedding (t-SNE) was used to jointly visualize cells/nuclei from all stages. Briefly, we obtained two-dimensional projections of cell population by first reducing the dimensionality of the gene expression matrix and then projecting these genes in to the t-SNE (van der Maaten & Hinton, 2008) space. We used Uniform Manifold Approximation and Projection (UMAP; McInnes et al., 2018) for visualization of cells/nuclei at their individual stages. UMAP was used for stage-specific embedding because it better preserved the global structure of the data and resulted in clearer depictions of the clusters at each time point. The following HDBSCAN settings were used to classify cells into clusters in an unbiased manner min_cluster_size = 6 and min_samples = 4.

### MARS clustering

We calculated differentially expressed genes among 42h APF ORN types that we identified in Li et al., 2020. We combined these genes with known antennal sensory receptors (Grabe et al., 2016), resulting in a total of 335 genes. We used this set of genes to confirm ORN types at 42h APF, and to identify ORN types at 24h APF and adult. To cluster each dataset, we applied MARS (Brbic et al., 2020)—a meta-learning approach for cell type discovery. MARS leverages annotations of previously annotated datasets to better separate cell types in an unannotated dataset. We first applied MARS to re-annotate 42h APF ORN types. The annotations agreed well with our previously identified types (Li et al., 2020), and we used them to annotate 24h APF. We ran MARS seven times with different random initializations (compared to 9 times on 42h APF transcriptomes) and neural network architectures to increase our confidence in the discovered clusters, and combined annotations from the different runs. A cluster was approved by calculating differentially expressed genes in that cluster and checking whether genes agree with known markers and/or are expressed uniquely in that cluster compared to other cells not assigned to the cluster. After annotating the 24h APF data, we used these annotations in combination with 42 APF annotations to guide the clustering of adult neurons. For adult annotations, we proceeded in the same way as when annotating 24h APF and ran MARS ten times followed by validating each cluster using marker genes.

### Transcriptome similarity analysis

To compare the transcriptome similarity between 24h APF, 42h APF, and adult clusters, we computed a set of unbiased 416 differentially expressed genes across all three developmental stages. For each cluster, we calculated expression profile of differentially expressed genes by taking the average across all cells within that cluster. We then calculated Pearson correlation coefficient of selected genes between all pairs of MARS clusters at each stage. For the transcriptome similarity analysis of all ORN types (Figure 4—supplement 2A), we used all clusters discovered by MARS. For the transcriptome similarity analysis within sensilla types (Figure 7E–G), we used only MARS clusters that were annotated and matched across all stages to guarantee consistent set of clusters for the comparison.

### Gene Ontology and STRING analysis

Gene Ontology (GO) analysis was used to characterize transcriptome changes in multiple cases. We generated gene lists for GO analysis by comparing the two groups using differential expression analysis (Mann-Whitney U test). We identified roughly equal numbers of differentially expressed genes that reached a significance level of p < 10^−5^ (after Bonferroni adjustment for multiple testing) for each comparison group and uploaded these gene lists into Flymine for GO analysis (Lyne et al., 2007). Redundant GO terms were removed using REVIGO (Supek et al., 2011). We report the p-value of enrichment for each term.

The STRING network was made using the top 75 differentially expressed genes from 24h APF ORN nuclei and adult nuclei into the STRING database search portal. Nodes that did not connect to the network were not displayed, and edges (links between genes) were generated by their reported interactions and corresponding confidence scores and plotted in Cytoscape (v3.7.1). Clusters were labeled using the GO terms that they were associated with and colored based on their on their fold change which was calculated by log_2_[(mean_adult_) / (mean_24h APF_ + 0.1)].

### Generation of dendrograms for 24h APF and adult transcriptomes

Dendrograms for 24h APF and adult data were derived from hierarchical clustering of annotated clusters at each stage. Pair-wise comparisons between all annotated clusters at each stage were performed to generate a list of differentially expressed genes. This resulted in 437 differentially expressed genes at 24h APF and 936 differentially expressed genes in adult clusters. Average gene expression was then computed for each cluster and differentially expressed genes were used for dimensionality reduction at their respective stages, followed by hierarchical clustering using the cluster map function of seaborn based on the Euclidean distance between each cluster.

### Manual (marker-based) matching and annotating of ORN types

ORN clusters were manually matched using a combination of differentially expressed marker genes and olfactory receptor (Ir, Gr, Or) expression. Marker genes were identified using our previously published cluster-specific genes at 42h APF (Li et al., 2020) or via a Mann-Whitney U test to find genes that are highly expressed in one (or a few) clusters compared to the rest. We searched for either a single gene or a set of genes whose expression was unique to a single cluster at both 24h APF and 42h APF. If these marker genes were expressed in one cluster at both time points, we considered these clusters to be the same ORN type. We used olfactory receptor expression to match some clusters from 24h to 42h APF. Olfactory receptors were exclusively used to annotate adult clusters and to match them to their corresponding 42h cluster. Genes used to annotate, and match clusters are summarized in dot plots in Figures 4 and 5.

### Automatic matching of ORN types

To automatically identify the same ORN types across different developmental stages, we first calculated differentially expressed genes in MARS clusters at each stage. We started by matching clusters between 24h and 42h APF, then we matched clusters between 42h and adult APF, and finally used matching between adult and 24h APF to confirm matches. To match clusters between two stages, we computed the similarity score of differentially expressed genes between all pairs of clusters using Jaccard similarity index, defined as the ratio of the number of elements in the intersection of two gene sets and the number of elements in the union of the gene sets. Two-way matches occurred when cluster X in one stage had the highest similarity to cluster Y in another stage, and the opposite also held true where cluster Y has the highest similarity to cluster X. One-way matches occurred when cluster X at one stage has the highest similarity to cluster Y at another stage, but cluster Y is most similar to another cluster. Since each cluster has a one-way match by design of the approach, we relied on one-way matching exclusively for those clusters whose transcriptomes are intermingled (inseparable) between 24h and 42h APF (Figure 7—supplement 1A).

### Quantification and statistical analysis

All RNA-seq data analysis was performed in Python using NumPy, SciPy, seaborn, pandas, scikit-learn, Scanpy (Wolf et al., 2018) and custom scRNA-seq analysis modules (Li et al., 2017; Brbic et al., 2020).

## Acknowledgements

We are grateful to Daniel Pederick, Jan Lui, Chuanyun Xu, David Luginbuhl, and Zhuoran Li for helpful discussions about this work and comments on the manuscript. We thank all members of the Luo lab for helpful discussions. We thank the Bloomington *Drosophila* Stock Center, the Vienna *Drosophila* Resource center, and the Developmental Studies Hybridoma Bank for fly lines and antibodies. We are grateful to Mary Molacavage for administrative support. Finally, we thank James Ferguson for help making the diagram in Figure 1A.

## Additional information

### Author contributions

Colleen McLaughlin, Conceptualization, Methodology, Software, Validation, Resources, Formal Analysis, Investigation, Writing–original draft preparation, Data curation, Writing–review & editing; Visualization. Maria Brbić, Methodology, Software, Formal Analysis, Data curation, Writing–review & editing, Investigation, Visualization. Qijing Xie, Investigation. Tongchao Li, Investigation. Felix Horns, Investigation, Resources. Sai Saroja Kolluru, Resources. Justus M. Kebschull, Resources. David Vacek, Investigation. Anthony Xie, Investigation. Jiefu Li, Investigation. Robert C. Jones, Resources. Jure Leskovec, Resources. Steven R. Quake, Resources, Funding Acquisition. Liqun Luo, Conceptualization, Resources, Writing–original draft, Writing–review & editing, Supervision, Funding Acquisition. Hongjie Li, Conceptualization, Methodology, Formal Analysis, Software, Investigation, Resources, Data Curation, Writing–original draft, Supervision.

---

**Figure 2—supplement 1.**
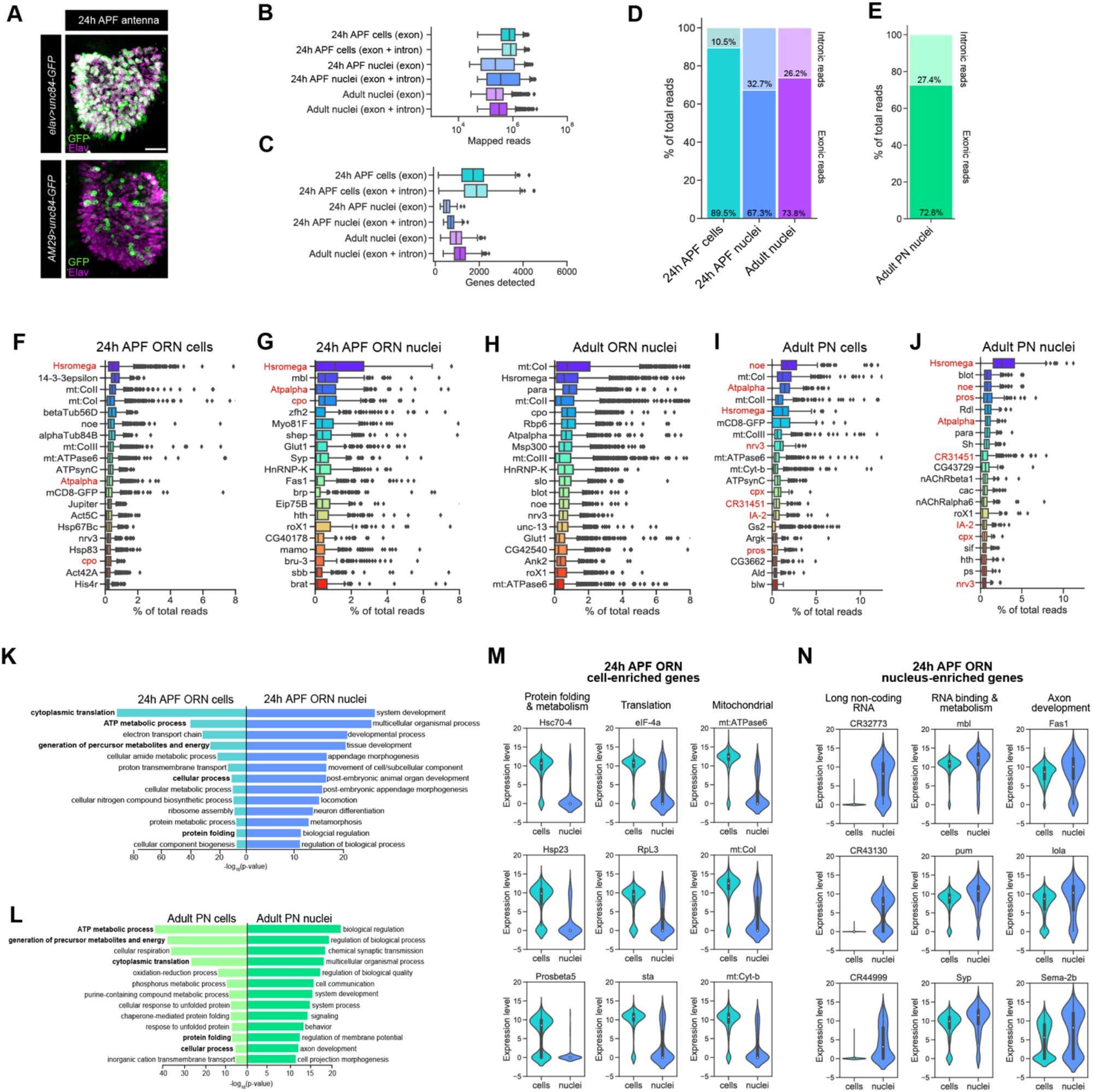
Additional comparisons of scRNA-seq and snRNA-seq protocols. (**A**) Confocal images showing expression of the nuclear envelop protein Unc84-GFP in the antenna driven by *elav-GAL4* and *AM29-GAL4*; anti-GFP labeling is in green and anti-Elav (pan-neuronal nuclear marker) labeling is in magenta. Scale bar, 20 μm. (**B–C**) Box plots showing distribution of uniquely-mapped reads (B) and genes detected (C) per cell/nucleus in ORNs. These plots depict a breakdown of all cells/nuclei with exonic reads only and exonic and intronic reads combined. Analyses in Figure 2 and the rest of the study were performed on cells with exonic reads only and nuclei with combined exonic and intronic reads. (**D–E**) Stacked bar plots of proportion of exonic and intronic reads in scRNA-seq and snRNA-seq samples where reads were combined in ORNs (D) and PNs (E). (**F–J**) Top 20 transcripts with highest percentage of total reads in 24h APF ORN cells (F), 24h APF ORN nuclei (G), adult ORN nuclei (H), adult PN cells (I), and adult PN nuclei (J). Note that top transcripts in adult ORN nuclei contain mitochondrial (mt) transcripts, likely artifacts resulting from dissociation procedures. Transcripts shared between stage-matched cells and nuclei are in red. (**K–L**) Gene ontology (GO) analysis based on genes enriched in 24h APF ORN cells compared to 24h APF ORN nuclei (K) and adult PN cells compared to adult PN nuclei (L). The top 13 significant GO terms are shown. ~500 DE genes were used for GO analysis in each category. Bolded terms in the cell categories are shared between ORN and PN cells. (**M–N**). Violin plots showing cell-enriched (M) and nucleus-enriched (N) transcripts in each category in 24h APF ORNs. Note that mitochondrial genes are more enriched in cells, whereas long non-coding RNAs are more highly enriched in nuclei.

**Figure 3—supplement 1.**
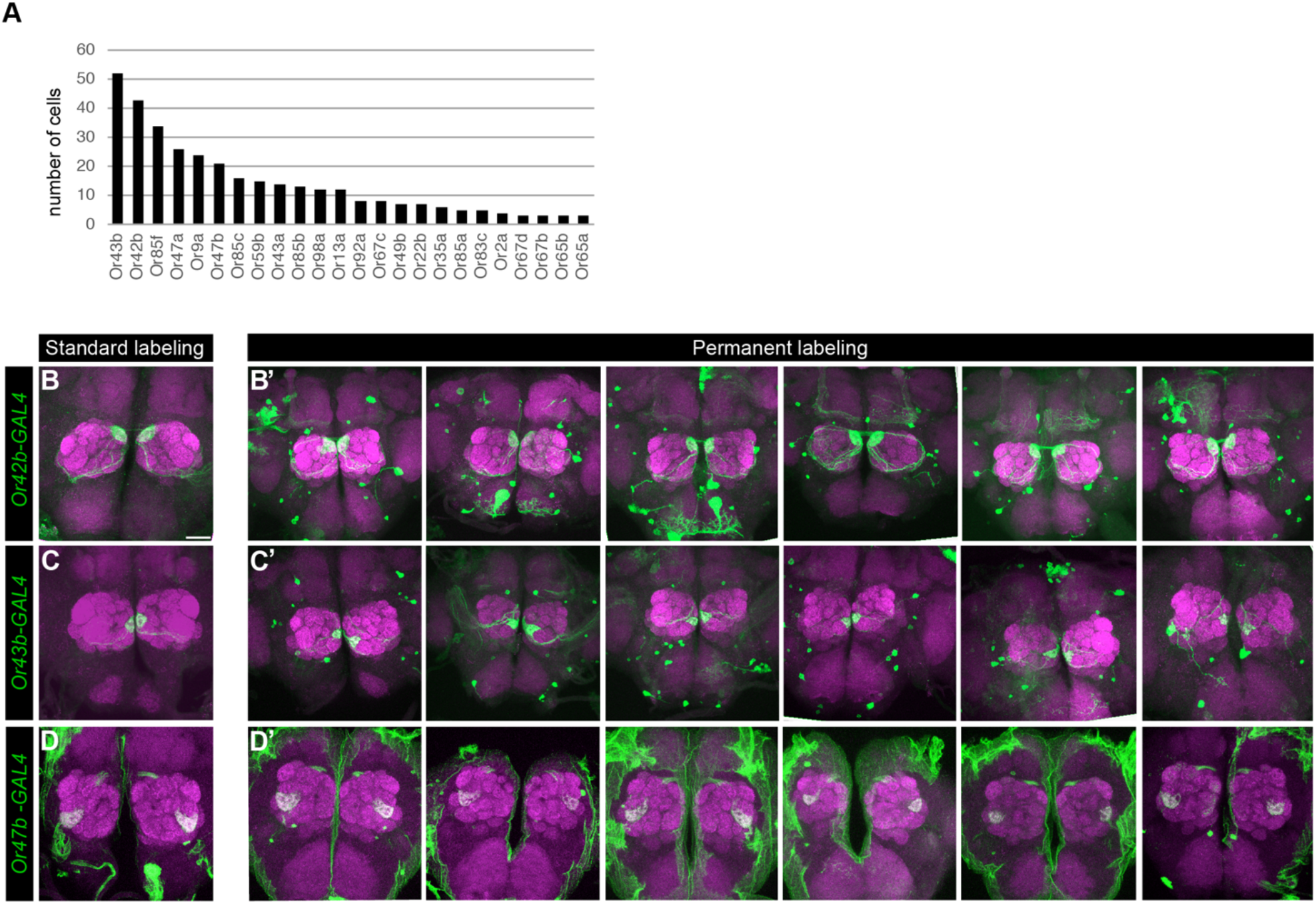
Permanent labeling reveals that *Drosophila* ORNs express the same receptor across all developmental stages. (**A**) Quantification of number of cells expressing each olfactory receptor in 42h APF ORNs. (**B–D’**) Additional confocal images of adult antennal lobes labeled with anti-GFP (green) and anti-NCad (magenta) in standard (B–D) and permanent labeling (B’–D’) of *Or42b-GAL4*-positive (B panels), *Or43b-GAL4-positive* (C panels), *Or47b-GAL4-positive* (D panels) ORNs. Scale bar, 20 μm.

**Figure 4—supplement 1.**
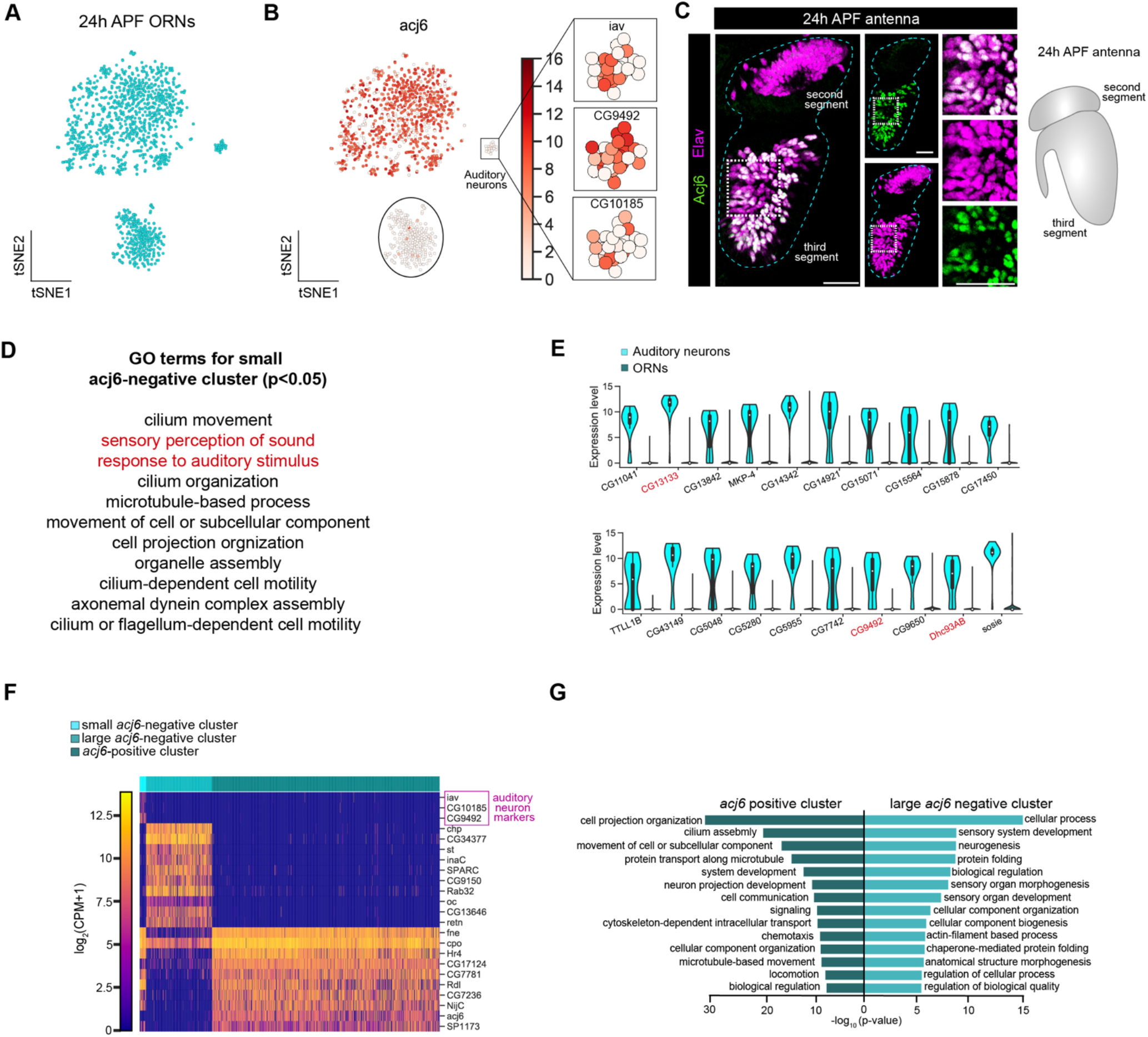
Two developmental states of 24h APF ORNs. (**A**) t-SNE plot showing that ORNs cluster into two large and one small population(s) of cells when dimensionality reduction was performed using the top 500 over-dispersed genes. (**B**) t-SNE plot of *acj6* in 24h APF neurons (left). Expression of three auditory neuron markers *iav, CG9492, CG10185* (right) in the small *acj6*-negative cluster. These markers are essentially absent from the two larger clusters in this plot. (**C**) Confocal images (left) of 24h APF antenna co-labeled with anti-Acj6 (green) and anti-Elav (magenta). Some cells are Elav-positive but Acj6-negative in the third antennal segment (zoomed-in panels to the right). White squares denote the location where zoomed in panels were obtained. Schematic of 24h APF antennal anatomy (far right). Scale bars, 20 μm. (**D**) List of GO terms enriched in the small *acj6*-negative cluster derived from the top 110 highest expressed genes in that cluster. Red terms are auditory organ-specific. (**E**) Violin plots of the top auditory-neuron enriched genes. Genes in red have known expression or function in auditory organs. (**F**) Heatmap depicting auditory neuron markers (top 3 genes; marked with magenta); top 10 DE genes between *acj6+* and large *acj6*-cluster (middle 10 genes), and the top 10 highest expressed genes in the *acj6+* cluster compared to the large *acj6*− cluster (last 10 genes). (**G**) Gene ontology (GO) analysis based on > 500 differentially expressed genes between *acj6+* and the large *acj6*− cluster. Top 14 significant GO terms are shown.

**Figure 4—supplement 2.**
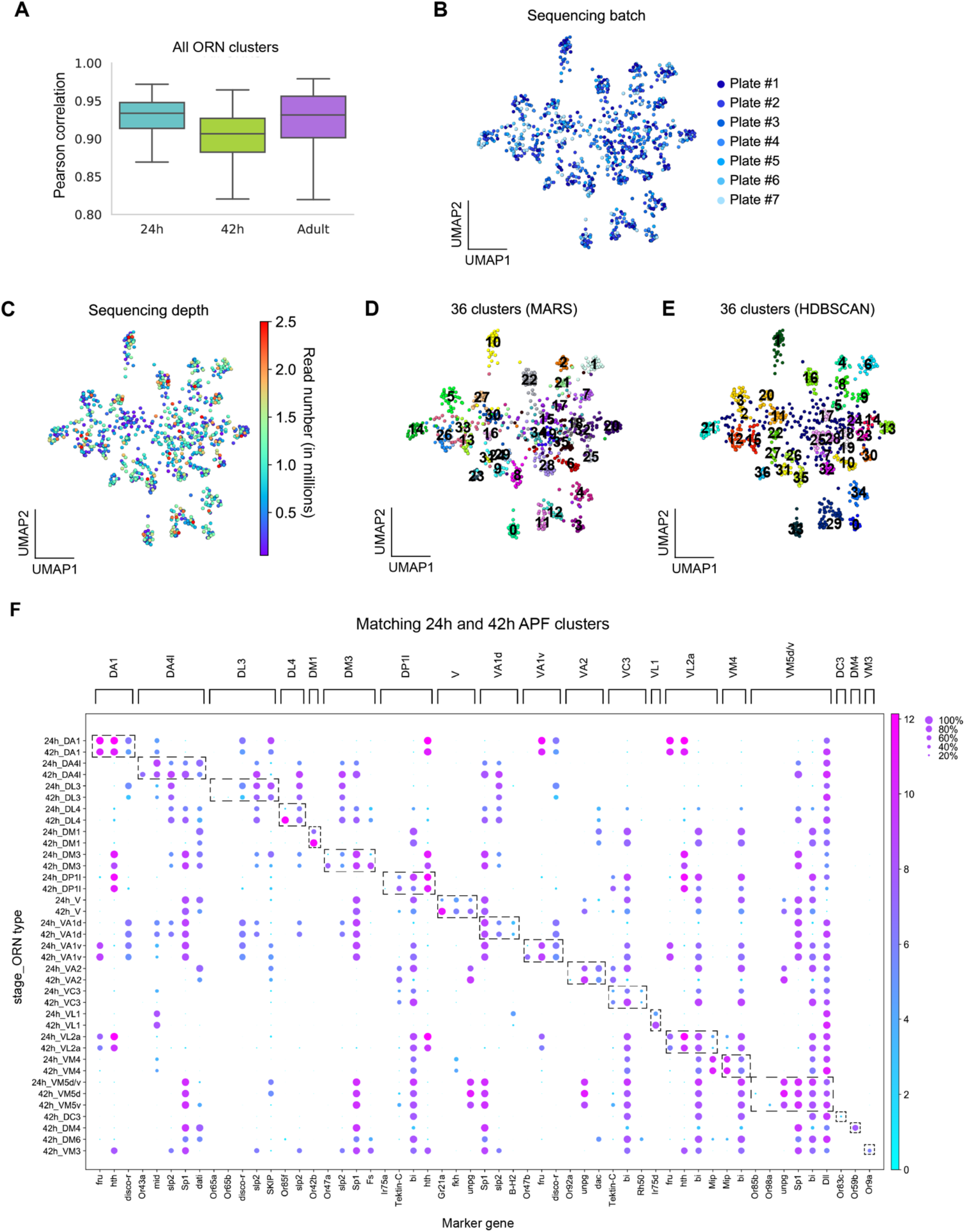
Quality control and additional analysis of 24h APF transcriptomes. (**A**) Quantification of cluster-level transcriptomic similarity of all ORN clusters annotated in this study and in Li et al., 2020 using Pearson’s correlation. 416 differentially expressed genes identified from all three stages were used for this analysis. (**B–C**) UMAP plots depicting 24h APF neurons color-coded by sequencing batch (plate #; B) and colored with a heat map depicting read number (C). (**D–E**) UMAP plots comparing MARS clustering (D) with HDBSCAN density-based clustering (E) approaches. MARS is better able to separate VA1v and VA1d [clusters # 11 and 12 in (D) and cluster # 29 in (E)]. (**F**) Dot plot showing expression of the markers used to match 24h APF ORNs to 42h APF ORNs. Some 42h APF clusters express olfactory receptors that are not found in 24h APF clusters. Receptors/markers of DM6 ORNs are not shown because sequencing of a cells labeled with a GAL4 driver was used to decode this cluster at 42h APF (Li et al., 2020). Dashed boxes highlight the markers for each cluster. Each dot represents: mean expression within each cluster (color) and fraction of cells within the cluster expressing the gene (dot size). Note this plot also depicts 42h APF DC3, DM4, DM6, and VM3 clusters as they were decoded in Li et al., 2020 but we were not able to annotate these clusters until Figure 6.

**Figure 4—supplement 3.**
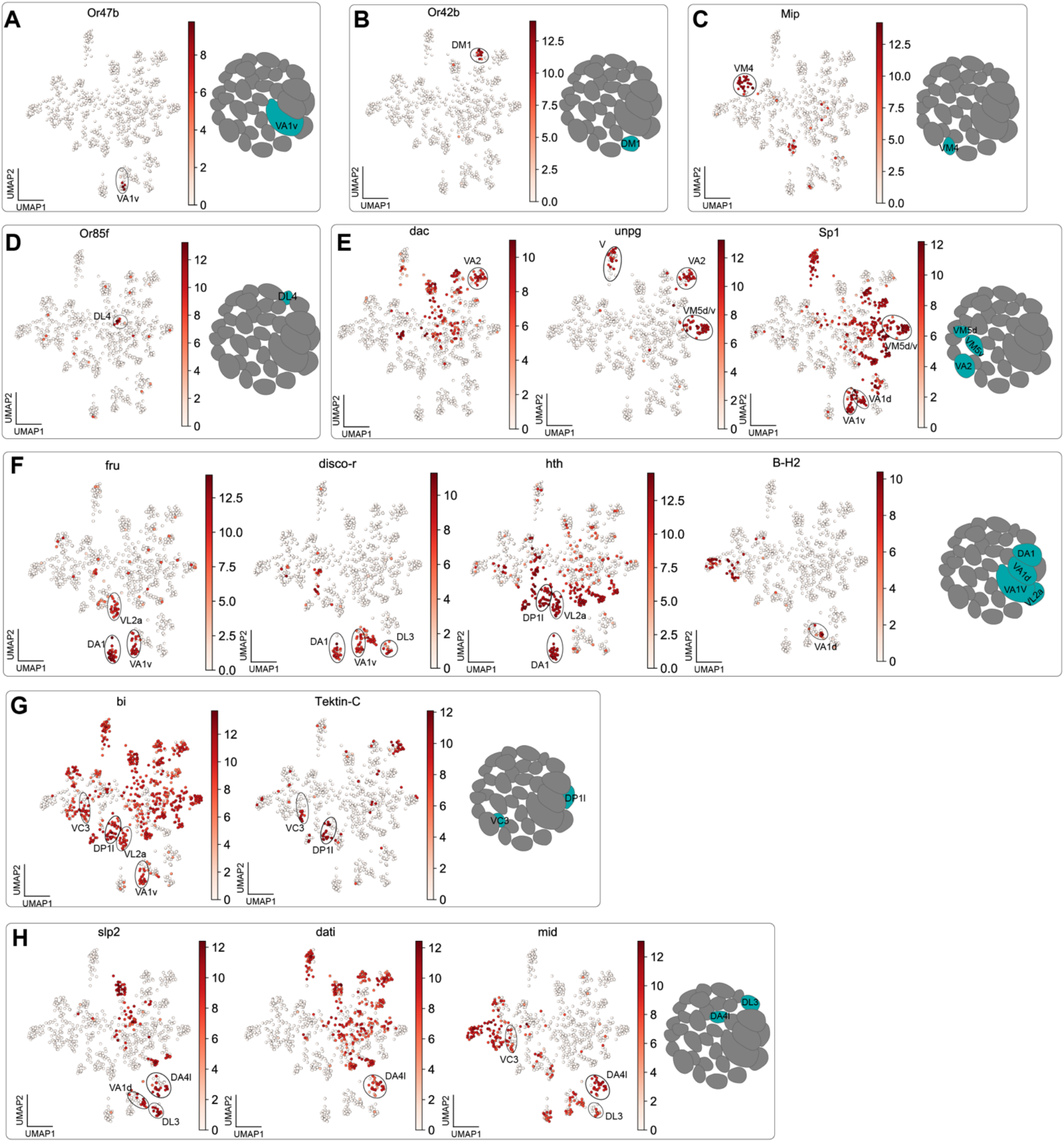
Markers used to match 24h APF antennal neurons to their glomerular type. (**A–H**) In each panel, a UMAP plot showing the gene(s) used to annotate clusters is next to a schematic depicting the glomerular target(s) of annotated neuron type(s). Heatmap scale bars are in log_2_(CPM + 1).

**Figure 5—supplement 1.**
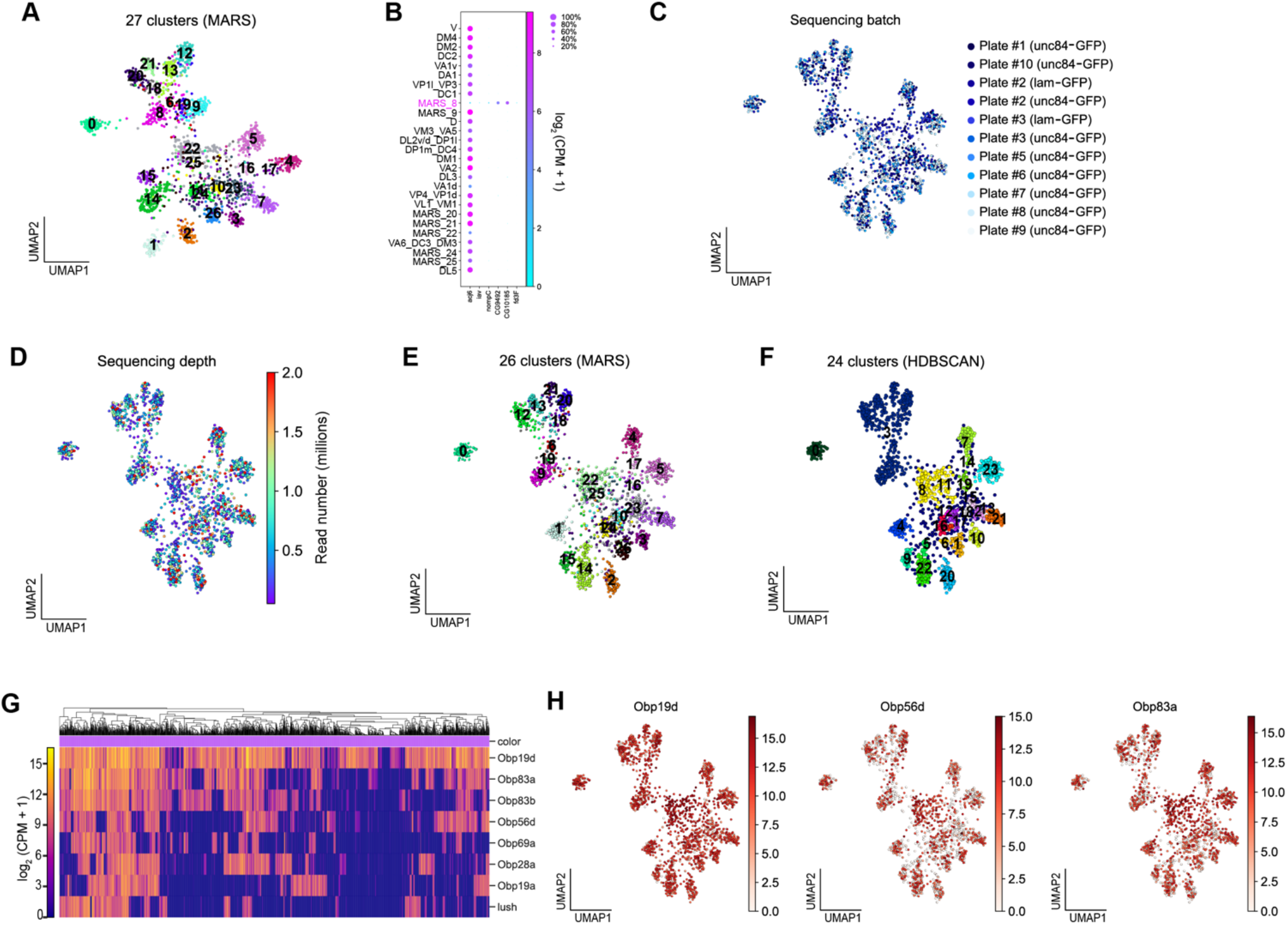
Quality control and additional analysis of adult transcriptomes. (**A–B**) UMAP plot of 27 clusters of adult neurons; note that MARS_8 was removed from subsequent analysis because it was found to express auditory neuron genes and not the ORN gene, *acj6*, shown in the dot plot (B). (**C–D**) UMAP plots depicting adult transcriptomes color coded by sequencing batch (plate #; C) and sequencing depth (D). Note that we included some adult nuclei labeled with lam-GFP to increase cell type representation in the adult data. (**E–F**) UMAP plots comparing MARS (E) with HDBSCAN density-based clustering (F). Both algorithms yield mostly concordant results for Or-expressing clusters; however, MARS predictions are better able to separate Ir-expressing clusters [compare top clusters in (E) with the same clusters in (F)]. (**G–H**) Heatmap depicting odorant binding protein (*Obp*) expression (G) and UMAP plots of select *Obps* showing that these genes are broadly expressed in adult neurons (H). Heatmap scale bars are in log_2_(CPM + 1).

**Figure 5—supplement 2.**
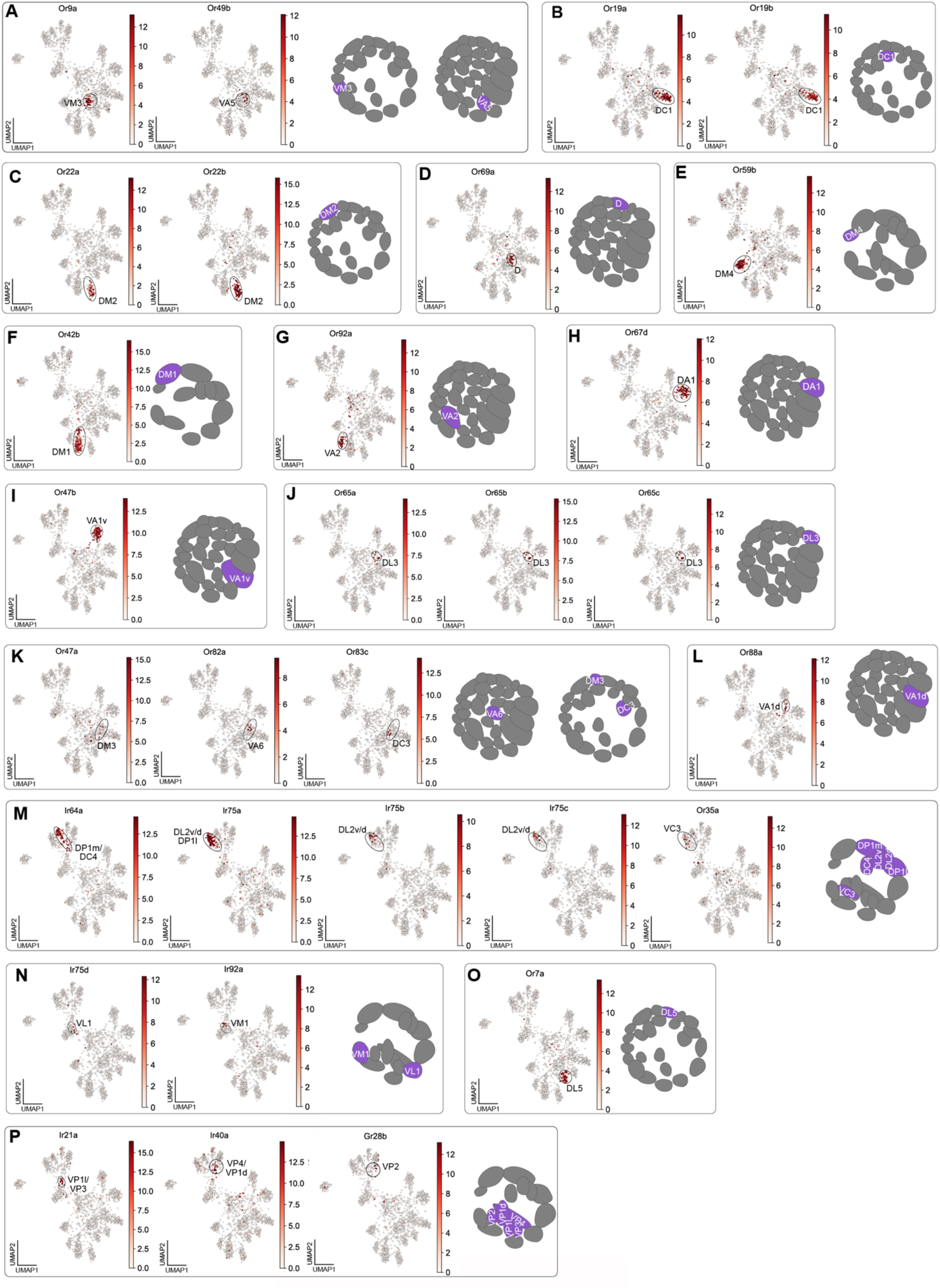
Cluster-specific sensory receptor expression in adult transcriptomes. (**A–P**) UMAP plots of sensory receptors used to annotate adult sensory neuron clusters. In each panel, a UMAP plot showing the gene(s) used to annotate clusters is placed next to a schematic showing the glomerular target(s) of decoded neuron type(s). Heatmap scale bars are in log_2_ (CPM + 1).

**Figure 6—supplement 1.**
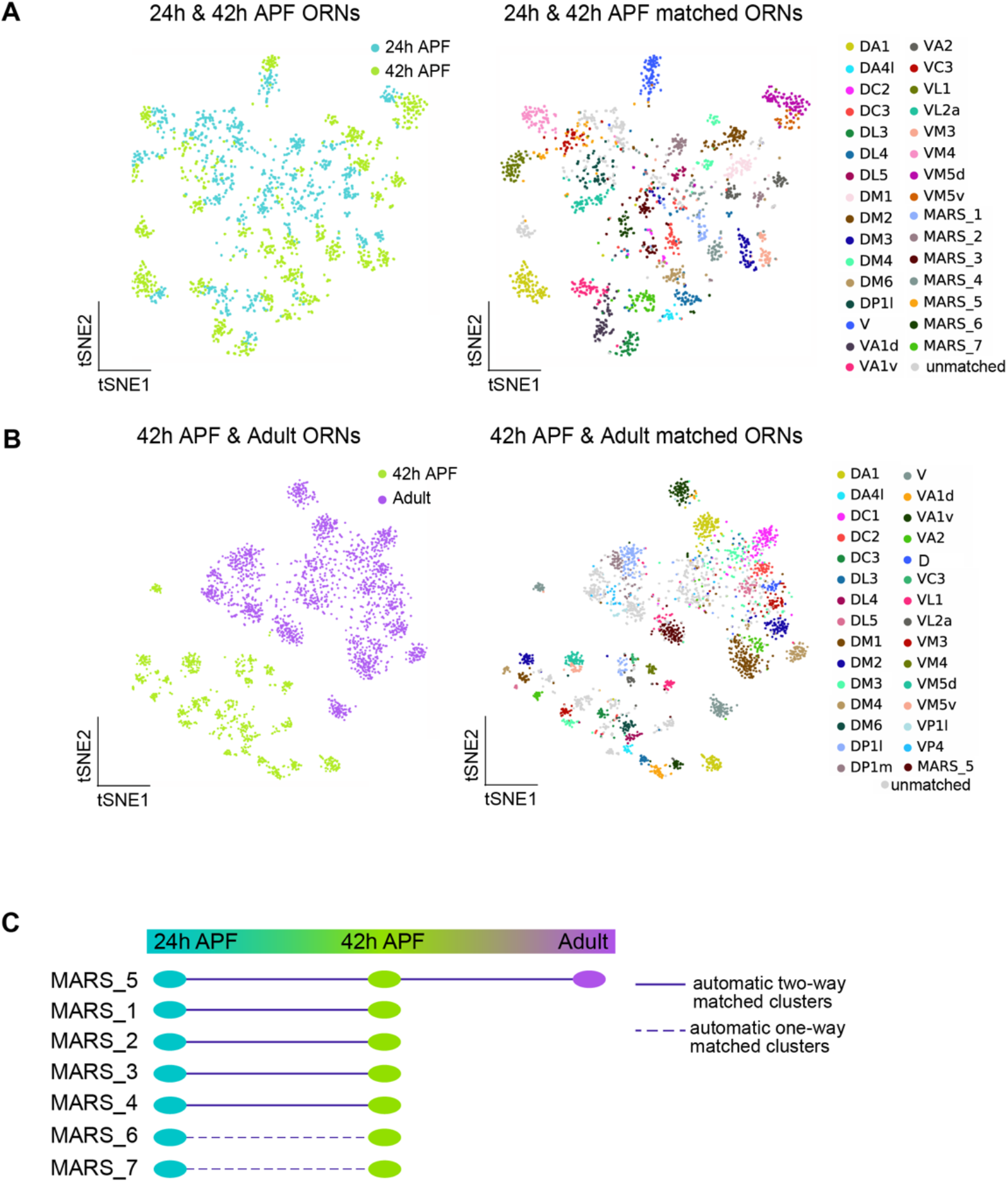
Additional evidence for cluster matching across developmental stages. (**A**) t-SNE plots depicting 24h APF and 42h APF ORNs (left) and matched clusters between these two timepoints (right). Dimensionality reduction was performed using differentially expressed genes from 42h APF combined with olfactory receptors for a total of 335 genes. “MARS_#” are matched between both timepoints but their glomerular identity is unknown. (**B**) t-SNE plots depicting 42h APF and adult ORNs (left) and matched clusters between these two timepoints (right). Dimensionality reduction was performed using differentially expressed genes from 42h APF combined with olfactory receptors for a total of 335 genes. MARS_5 is matched across all three timepoints but its glomerular identity is unknown. (**C**) Summary of matched ORN types where the glomerular identity of these clusters is unknown.

**Figure 7—supplement 1.**
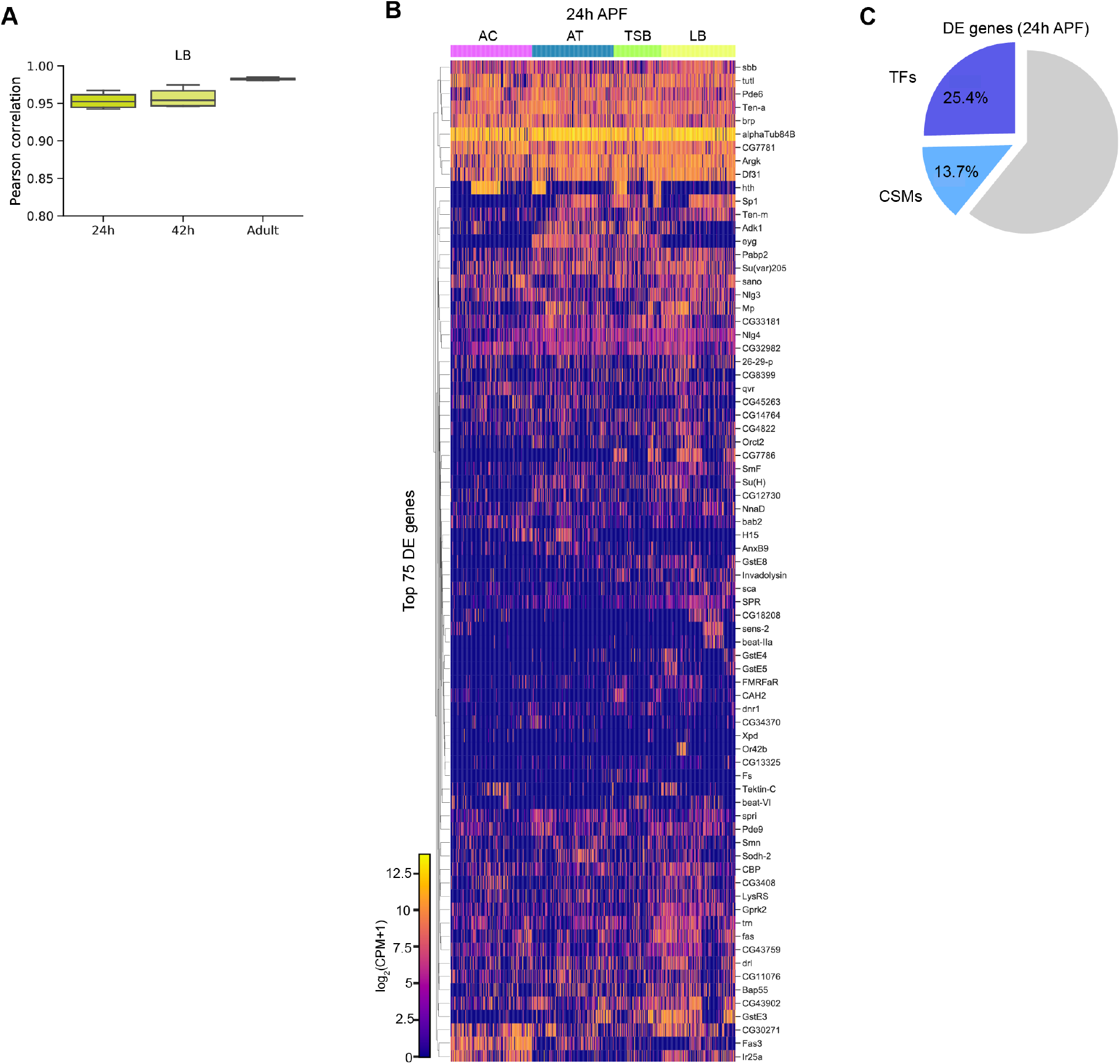
Comparisons of 24h APF ORNs by sensillar class. (**A**) Quantification of cluster-level similarity of ORNs within the large basiconic sensillar class excluding V neurons, at all stages profiled. (**B**) Heatmap of top 75 differentially expressed genes in 24h APF ORNs grouped by sensillar class. (**C**) Quantification of the percentage of TFs and CSMs in the 437 differentially expressed genes used to cluster 24h APF transcriptomes in (Figure 7I).

**Figure 7—supplement 2.**
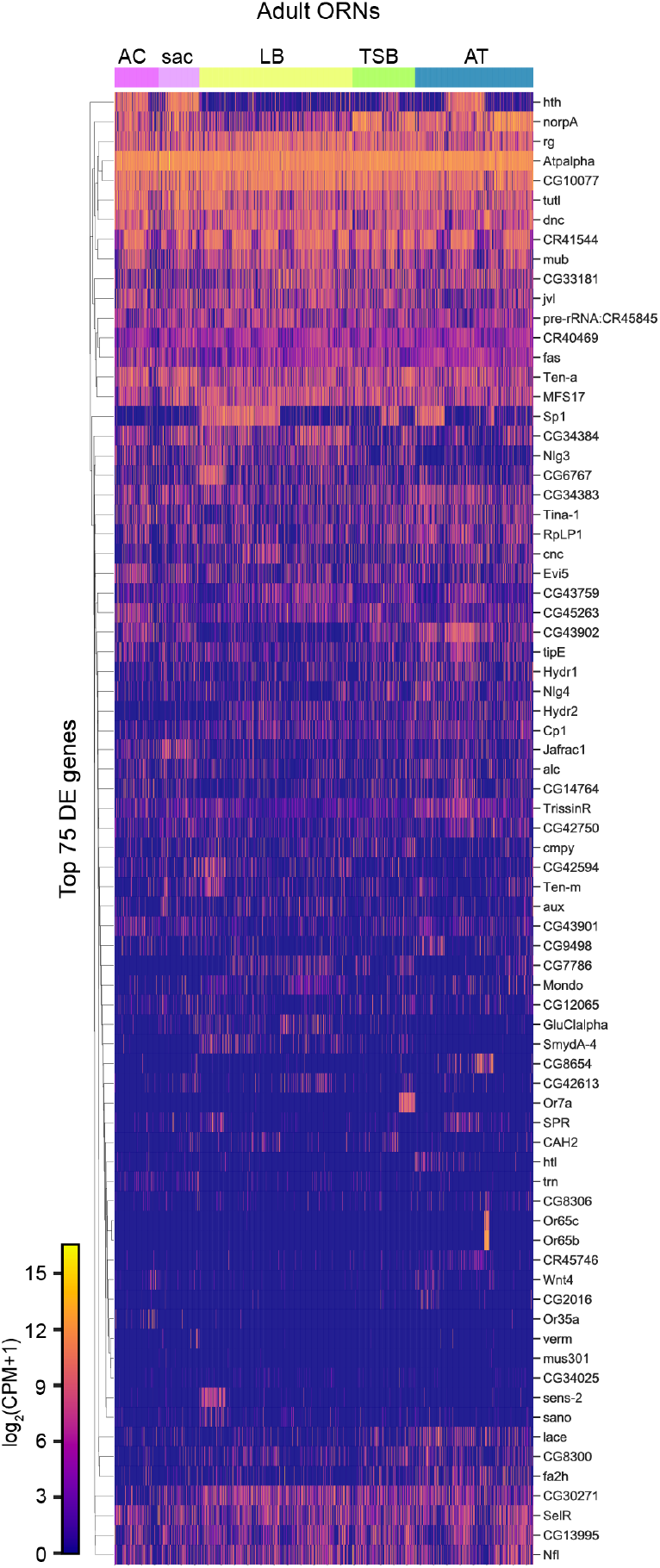
Comparisons of adult neurons by sensillar class and sensory structure. Heatmap of the top 75 differentially expressed genes in adult neurons grouped by sensory structure.

**Figure 8—supplement 1.**
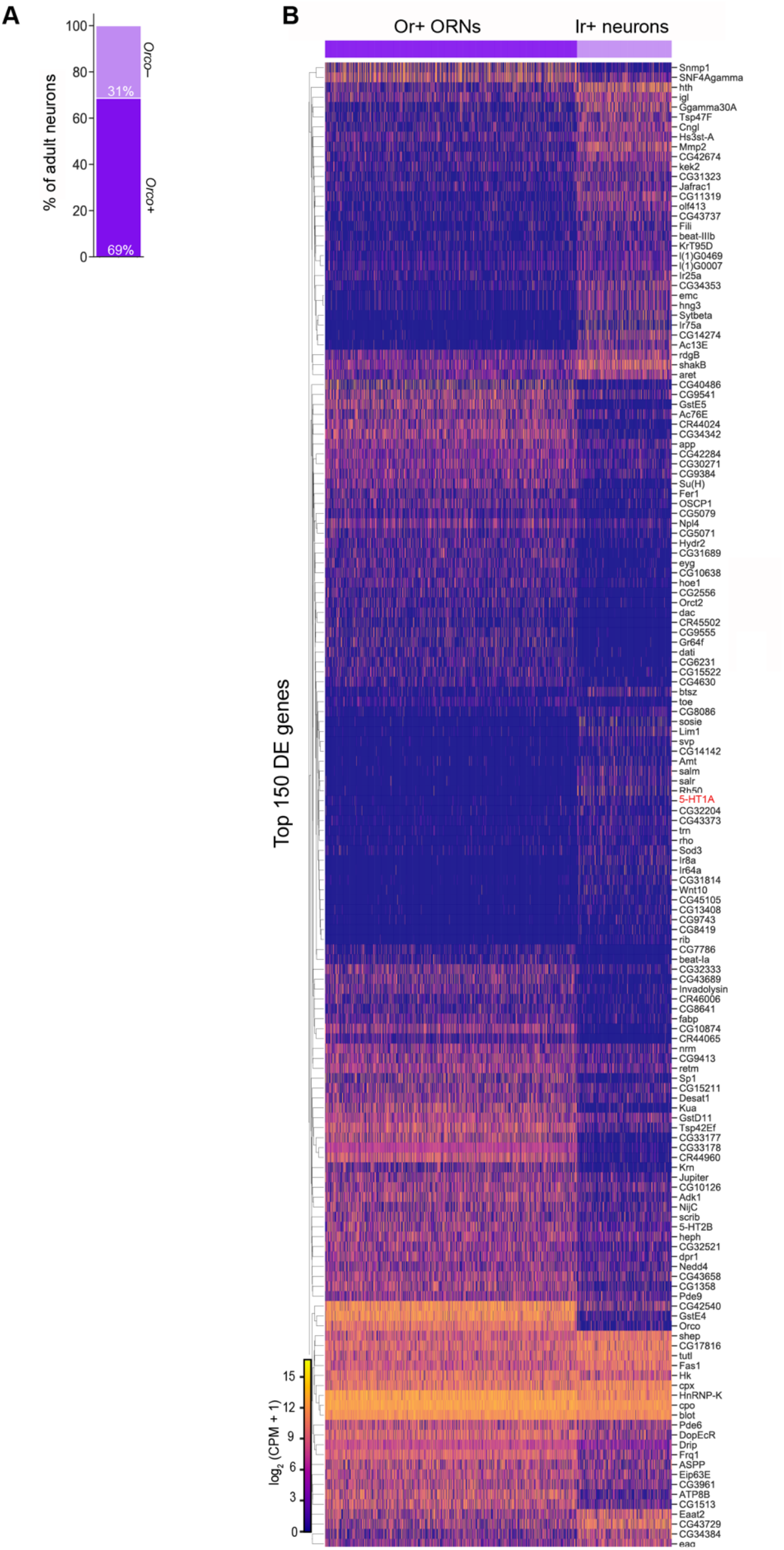
Differentially expressed genes in Or- and Ir-expressing neurons. (**A**) Stacked bar plot quantifying the percentage of adult antennal neurons that are *Orco* high (e.g., express *Orco* at log_2_(CPM+1) ≥ 5) and *Orco* low. (**B**) Heatmap of top 150 differentially expressed genes between Or and Ir clusters. *5-HT1A* expression is in red.

